# Juxtacrine DLL4-NOTCH1 signaling between astrocytes drives neuroinflammation via the IL-6-STAT3 axis

**DOI:** 10.1101/2023.10.04.560826

**Authors:** Pierre Mora, Margaux Laisné, Célia Bourguignon, Paul Rouault, Alain-Pierre Gadeau, Marie-Ange Renault, Sam Horng, Thierry Couffinhal, Candice Chapouly

## Abstract

Under neuroinflammatory conditions, astrocytes acquire a reactive phenotype that drives acute inflammatory injury as well as chronic neurodegeneration. We hypothesized that astrocytic DLL4 may interact with its receptor NOTCH1 on neighboring astrocytes to regulate astrogliosis via downstream juxtacrine signaling pathways. Here we investigated the role of astrocytic DLL4 on neurovascular unit homeostasis under neuroinflammatory conditions. We probed for downstream effectors of the DLL4-NOTCH1 axis and targeted these for therapy in two models of CNS inflammatory disease. We first demonstrated that astrocytic DLL4 is upregulated during neuroinflammation, both in mice and humans, driving astrogliosis and subsequent blood brain barrier permeability and inflammatory infiltration. We then showed that the DLL4-mediated NOTCH1 signaling in astrocytes directly drives IL-6 levels, induces STAT3 phosphorylation promoting upregulation of astrocyte reactivity markers, pro-permeability factor secretion and consequent blood brain barrier destabilization. Finally we revealed that blocking DLL4 with antibodies improves experimental autoimmune encephalomyelitis symptoms in mice, identifying a potential novel therapeutic strategy for CNS autoimmune demyelinating disease. As a general conclusion, this study demonstrates that DLL4-NOTCH1 signaling is not only a key pathway in vascular development and angiogenesis, but also in the control of astrogliosis during neuroinflammation.

## Introduction

Proper function of the healthy central nervous system (CNS) requires the selective regulation of soluble factors and immune cells through the blood–brain barrier (1). Blood brain barrier structure and integrity involves a complex network of crosstalk among various components of the neurovascular unit, including vascular endothelial cells, pericytes, microglia and the astrocyte endfeet (Glia limitans) (2).

Substantial intercellular communication occurs between endothelial cells and the astrocyte endfeet of the blood brain barrier (1, 3). We recently identified a capacity for bidirectional signaling between endothelial cells and astrocytes in the neurovascular unit (4). How these signals contribute to cerebrovascular impairment, blood brain barrier dysfunction and neuroinflammation remains unclear and carries translational implications for autoimmune demyelinating diseases, stroke and other CNS inflammatory conditions.

Under neuroinflammatory conditions, astrocytes acquire a reactive phenotype, defined by morphological and molecular changes that drive acute inflammatory injury as well as chronic neurodegeneration. This “reactive astrogliosis” occurs in a context-specific manner with specific pathways and phenotypes recruited in response to the type, severity and timing of CNS injury. Furthermore, simultaneous induction of both pathogenic and protective pathways may be induced. For instance, during infections (HIV, Herpes virus), and in Alzheimer’s disease, multiple sclerosis and Parkinson’s disease, reactive astrocytes produce both pro-inflammatory factors such as endothelin1, glutamate, interleukin1β, Tumor Necrosis Factor and nitrite oxide and neuroprotective factors such as nerve growth factor and glial cell line-derived neurotrophic factors (5). We previously identified that reactive astrocytes induce blood brain barrier opening in inflammatory conditions via production of VEGFA and TYMP (6) which downregulate the endothelial tight junction proteins that protect the blood brain barrier in the healthy brain (7).

NOTCH1 receptor is a central effector of astrogliosis in a wide range of neuropathological contexts. After intra-cerebral hemorrhage (8), in stroke (9) and in several neuroinflammatory conditions such as amyotrophic lateral sclerosis (10), multiple sclerosis and experimental auto-immune encephalomyelitis (EAE) (11, 12), NOTCH1 signal transduction is activated in reactive astrocyte populations.

Notch signaling, which is highly conserved in vertebrates, is stimulated by the interaction of Notch receptor with its ligands, Delta and Jagged, trans-membrane proteins with large extracellular domains. Ligand binding promotes two proteolytic cleavages in the Notch receptor: the first catalysed by ADAM-family metalloproteases and the second mediated by γ-secretase (13–15). The second cleavage releases the Notch intracellular domain (NICD), which translocates to the nucleus and cooperates with the DNA-binding protein CSL (named after CBF1, Su(H) and LAG-1) to promote transcription. The precise numbers of NOTCH receptors differ between species (16): in vertebrates, 4 different NOTCH receptors have been identified and in the neurovascular unit, NOTCH1 is expressed by astrocytes and endothelial cells, NOTCH4 by endothelial cells (17–19) and NOTCH3 by mural cells, or pericytes (20, 21).

Interestingly, activation of the NOTCH1-STAT3 axis in EAE has been shown to control the production of inflammatory cytokines by reactive astrocytes via the long non-coding RNA Gm13568, which has also been implicated in MS pathogenesis. Moreover, the NOTCH1-STAT3 pathway has similarly been identified as an effector of inflammation-induced differentiation of neurotoxic A1 astrocytes in a model of spinal cord injury and glial scar formation (22, 23).

As opposed to the abundant literature on the role of NOTCH1 receptor, DLL4 ligand expression has only been reported once, anecdotally, in reactive astrocytes, following brain injury (24). The important role of DLL4-NOTCH1 signaling in the cardiovascular system is already widely appreciated, but little is known about this specific pathway in reactive astrogliosis during neuroinflammation. We hypothesized that astrocytic DLL4 may interact with the receptor NOTCH1 on neighboring astrocytes to regulate astrogliosis and neuroinflammation via downstream juxtacrine signaling pathways.

We first tested the role of astrocytic DLL4 on neurovascular unit homeostasis under neuroinflammatory conditions. We then probed for downstream effectors of the DLL4-NOTCH1 axis and targeted these for therapy in two models of CNS inflammatory disease. Here, we demonstrate that astrocytic DLL4 is upregulated during neuroinflammation, both in mice and humans, driving astrogliosis and subsequent blood brain barrier permeability and inflammatory soluble factor and immune cell infiltration. We then show that the DLL4-mediated NOTCH1 signaling in astrocytes directly drives IL-6 levels, induces STAT3 phosphorylation promoting upregulation of astrocyte reactivity markers, pro-permeability factor secretion and consequent blood brain barrier destabilization. Finally we reveal that blocking DLL4 with antibodies improves EAE symptoms in mice, identifying a potential novel therapeutic strategy for CNS autoimmune demyelinating disease.

In sum, we report here for the first time that the DLL4-NOTCH1 axis acts as a key driver of astrogliosis during neuroinflammation via upregulation of the IL-6-STAT3-TYMP/VEGFA signaling pathway, leading to disruption of the neurovascular unit, increased immune infiltration into the CNS parenchyma and worsened neuropathology. More generally, this study demonstrates that DLL4-NOTCH1 signaling is not only a key pathway in vascular development and angiogenesis, but also in the control of astrocytic reactivity during neuroinflammation.

## Results

### DLL4 is expressed by CNS astrocytes and up-regulated during chronic neuroinflammation

First, we showed that *DLL4* is strongly up-regulated in reactive astrocytes under inflammatory conditions *in vitro* using CNS normal human astrocytes (NA) from ScienCell treated with IL-1β (interleukin-1β), a central pro-inflammatory cytokine driving multiple sclerosis pathophysiology (Fig 1, A, C-D) and *in vivo* (Fig 1, E), in the neurovascular unit, using experimental autoimmune encephalomyelitis (EAE), a pre-clinical model of multiple sclerosis using MOG_35-55_ to induce chronic neuroinflammation in *C57BL/6* mice. For this experiment, isolated spinal cord micro-vessels underwent a digestion step followed by a CD45^+^ T cell depletion step to eliminate inflammatory cell infiltrates induced by EAE. *DLL4* upregulation in human reactive astrocytes *in vitro* is associated with upregulation of the astrogliosis marker *TYMP1* (thymidine phosphorylase) (Fig 1, B). *Dll4* upregulation in the neurovascular unit is associated with upregulation of Notch signaling activation markers *Hey1* (hairy/enhancer-of-split related with YRPW motif protein 1) and *Jag1* (jagged 1) (Fig 1, F-G). We then confirmed that DLL4 is up-regulated in reactive astrocytes *in vivo*, on spinal cord sections from EAE induced C57BL/6 adult mice (Fig 1, H) and on cortical active lesions from multiple sclerosis patients (Fig 1, I).

**Fig 1:**
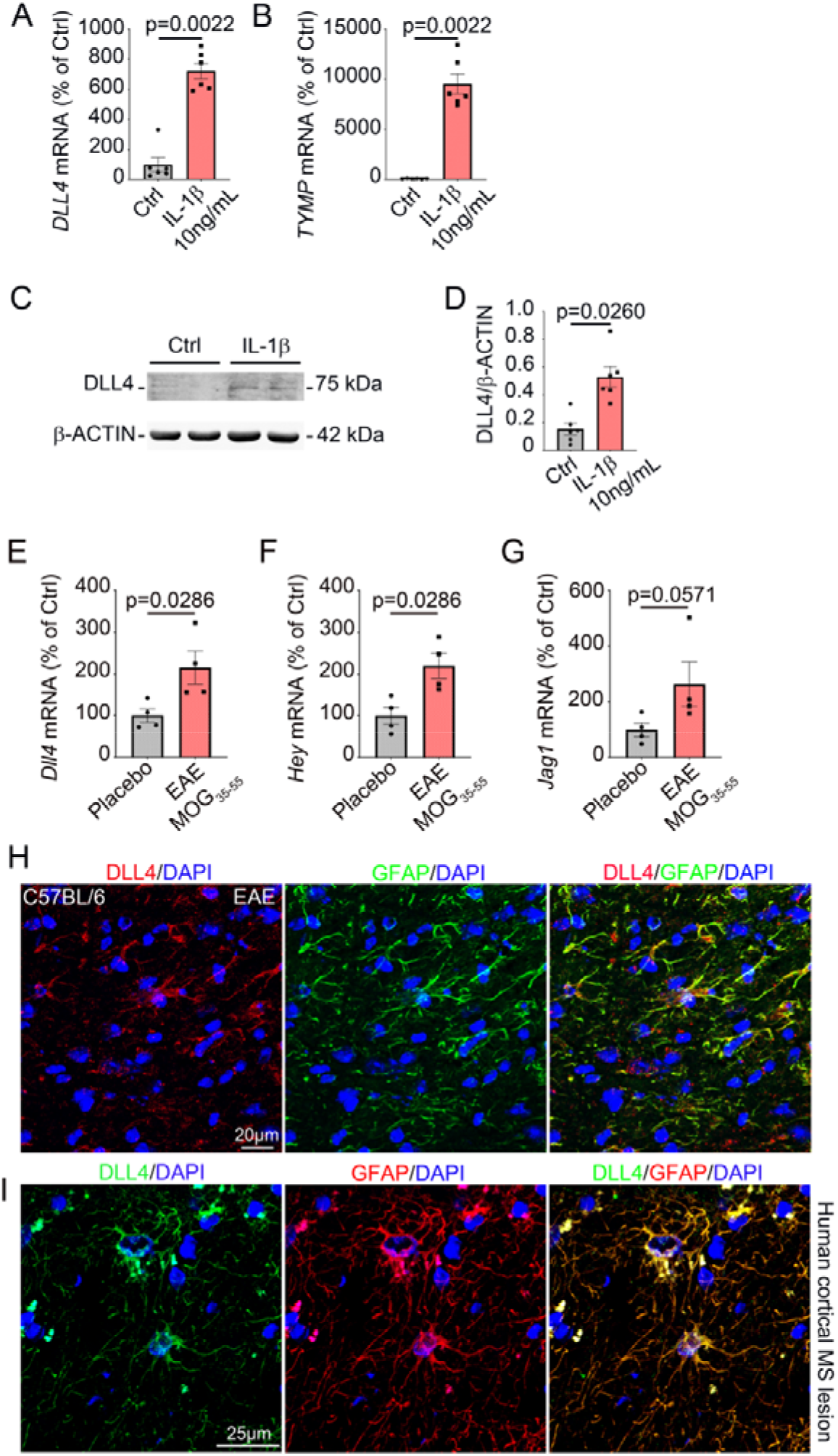
DLL4 is expressed by CNS astrocytes and up-regulated during chronic neuroinflammation: **(A-D)** Human Normal Astrocytes (NA) were seeded and cultured until confluence and starved for 24 h. NA were then treated for 24h with PBS 1X (control condition) versus 10 ng/mL of recombinant IL-1β (test condition). **(A)** *DLL4 and* **(B)** *TYMP* expression were then quantified by qRT-PCR. β*-ACTIN* was used as a reference. **(C)** Representative western blot for DLL4 and β-ACTIN protein expression level are shown. **(D)** DLL4 protein expression level was quantified by western blot. β-ACTIN was used as a reference. **(E-H)** 12-week-old *C57Bl/6* females (4 animals per group) were induced with MOG_35-55_ EAE versus Placebo. At day 13 post induction, mice were sacrificed and spinal cord neurovascular units were isolated. **(E)** *Dll4*, **(F)** *Hey1*, and **(G)** *Jag1* expression were measured via qRT-PCR in both groups (MOG_35-55_ versus placebo). **(H)** Spinal cord sections were harvested from MOG_35-55_ EAE induced versus Placebo treated *C57Bl/6* females and immune-stained with anti-DLL4 (in green) and anti-GFAP (in red) antibodies. Representative DLL4/GFAP staining is shown. **(I)** Human cortical sections from healthy donors and patients with cortical multiple sclerosis lesions were obtain from the NeuroCEB biobank and immunostained with anti-DLL4 (in green) and anti-GFAP (in red) antibodies. Nuclei were stained with DAPI (in blue). Representative DLL4/GFAP staining is shown. Mann-Whitney U test.

### Inactivation of astrocyte *Dll4* reduces disability in a model of multiple sclerosis during the onset and plateau of the disease

To test the role of astrocyte DLL4 in neuroinflammation, we conditionally disrupted *Dll4* expression in astrocytes using 2 different promoters (the *Glast-Cre^ERT2^* promoter and the *Aldh1L1-Cre^ERT2^* promoter) and examined the consequences on EAE pathology. Experimental mice consisted of *Glast-Cre^ERT2^, Dll4^Flox/Flox^* mice or *Aldh1L1-Cre^ERT2^, Dll4^Flox/Flox^* mice with corresponding littermate *controls (Dll4^Flox/Flox^*). We first verified the efficiency of both knockouts, 4 months after inducing knockdown by intra-peritoneal injection of tamoxifen. The *Glast-Cre^ERT2^*promoter induced a moderate recombination (30-50%) whereas the *Aldh1L1-Cre^ERT2^*promoter induced a full recombination (90-100%) in both spinal cord and cortical astrocytes (Supplemental Fig 1, A-H). The difference in recombination efficiency enabled us to test the effects of partial versus complete astrocyte *Dll4* knockdown in the disease course and spinal cord pathology of experimental multiple sclerosis (EAE).

In the rest of the manuscript, *Glast-Cre^ERT2^, Dll4^Flox/Flox^* mice will be named *Dll4^ACKO^P* (for partial recombination) and *Aldh1L1-Cre^ERT2^, Dll4^Flox/Flox^* mice will be named *Dll4^ACKO^C* (for complete recombination) to make reading easier.

We confirmed the absence of DLL4 astrocyte expression in non-reactive astrocytes in *Dll4^ACKO^C* mice versus *control* littermates (Supplemental Fig 1, I) and highlighted astrocyte specific DLL4 down-regulation in EAE induced *Dll4^ACKO^C* mice versus *control* littermates (Supplemental Fig 1, J) and in *Dll4^ACKO^P* mice versus *control* littermates injected in the cortex with AdIL-1β (Supplemental Fig 1, K)), another mouse model of astrogliosis and CNS autoinflammation.

In the *Dll4^ACKO^P* mouse model, *control* mice demonstrated neurologic deficits from day 11, which increased in severity until day 21, when the clinical score stabilized at a mean of 3.0, representing hind limb paralysis. In contrast, the clinical course in *Dll4^ACKO^P* mice was much milder: disease reached a plateau at day 27 at a mean of 2.5, indicating hind limb weakness and unsteady gait, a milder phenotype (Fig 2, A). The EAE peak score (Fig 2, B) was no different between the groups but average score during the time of disability (Fig 2, C) was decreased in *Dll4^ACKO^P* mice.

**Fig 2:**
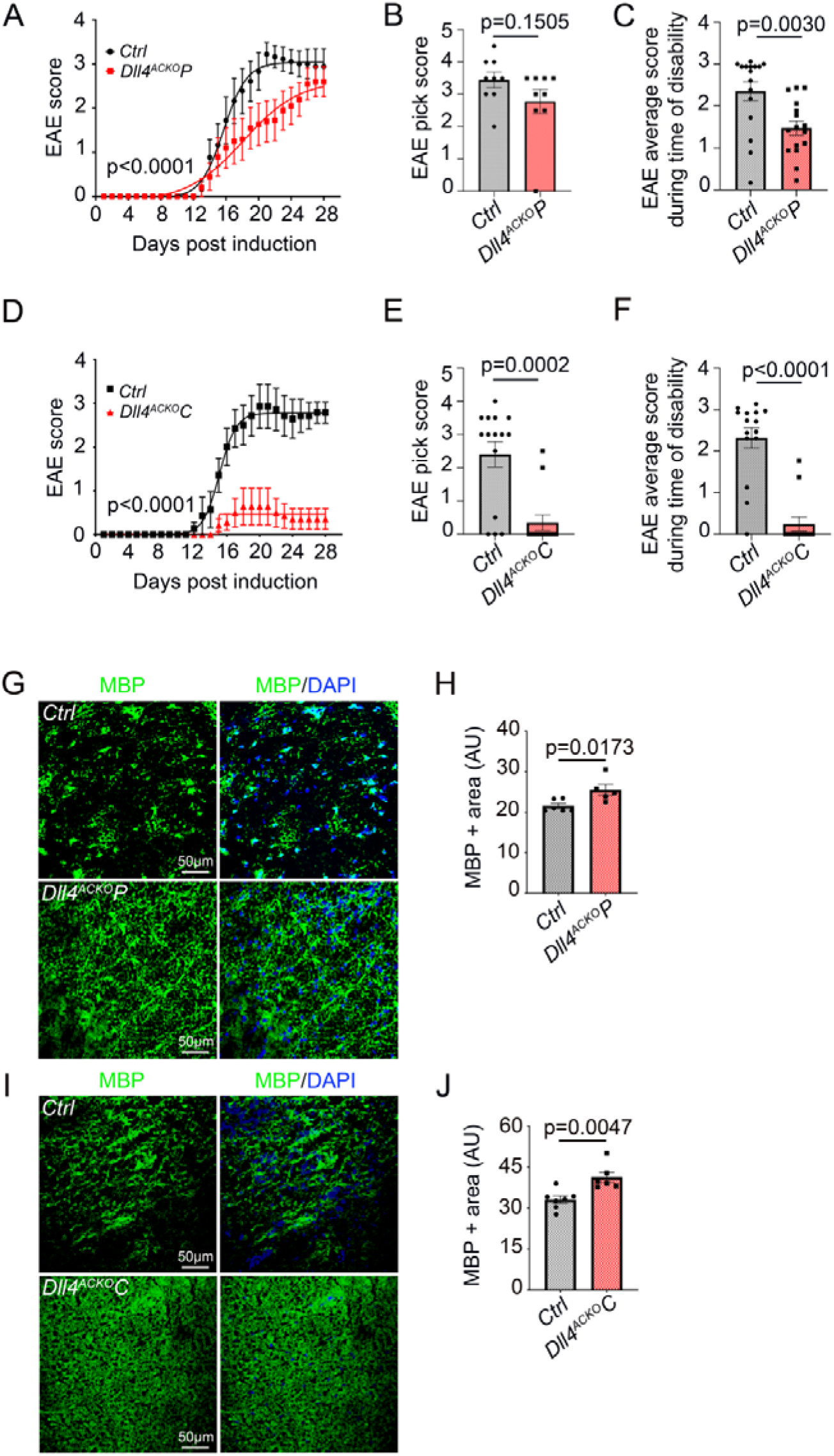
Inactivation of astrocyte *Dll4* reduces disability in a model of multiple sclerosis during the onset and plateau of the disease: **(A)** *Dll4^ACKO^P* mice and *control* mice induced with EAE were scored daily according to a widely-used 5-point scale (EAE scoring: 1 limp tail; 2 limp tail and weakness of hind limb; 3 limp tail and complete paralysis of hind legs; 4 limp tail, complete hind leg and partial front leg paralysis), nonlinear regression (Boltzmann sigmoidal). (*Dll4^ACKO^P n* = 10, WT *n* = 9). (B) *Dll4^ACKO^P* and *control* mice EAE peak score and **(C)** EAE average score during time of disability were quantified other the course of the disease. (D) *Dll4^ACKO^* mice and *control* mice induced with EAE were scored daily on a standard 5-point scale, nonlinear regression (Boltzmann sigmoidal). (*Dll4^ACKO^C n* = 15, WT *n* = 15). (E) *Dll4^ACKO^C*and *control* mice EAE peak score and **(F)** EAE average score during time of disability were quantified other the course of the disease. **(G-J)** Spinal cord EAE lesions from *Dll4^ACKO^P* mice, *Dll4^ACKO^C* mice and littermate *controls* were harvested at 18 days post induction. (G) *Dll4^ACKO^P* mice and *control* lesions and (I) *Dll4^ACKO^C* mice and *control* lesions were immune-stained with an anti-MBP (in green) antibody. Nuclei were stained with DAPI (in blue). **(H, J)** MBP positive areas were quantified **(H)** (*Dll4^ACKO^P* mice *n* = 5, WT *n* = 6), **(J)** (*Dll4^ACKO^C* mice n = 6, WT n = 7)). Mann-Whitney U test.

In the *Dll4^ACKO^C* mouse model, *control* mice exhibited neurologic deficits from day 12, which increased in severity until day 20, when clinical score stabilized at a mean of 2.8, representing hind limb paralysis. In contrast, the onset of clinical signs in *Dll4^ACKO^C* mice was first seen four days later, and the clinical course was very mild. Indeed, in *Dll4^ACKO^C* mice, disease reached a plateau at day 18 at a mean of 0.4 indicating almost no sign of paralysis (Fig 2, D). Strikingly, only two *Dll4^ACKO^C* mice developed very mild symptoms, the remaining majority showing no sign of pathology other the course of the disease. The EAE peak (Fig 2, E) and average scores during the time of disability (Fig 2, F) were both strongly decreased in *Dll4^ACKO^C* mice.

Clinical course in both *Dll4^ACKO^P* mice and *Dll4^ACKO^C* mice was correlated with decreased areas of demyelination as compared to the *control* cohorts (Fig 2, G-J).

In sum, we found that clinical course and pathology of EAE are reduced in mice with astrocyte *Dll4* knockdown. Furthermore, degree of *Dll4* knockdown appeared to have a dose-dependent effect as reduced pathology was only observed during the onset of the disease in the *Dll4^ACKO^P* mice while reduced pathology was observed during both the onset and plateau of the disease in the *Dll4^ACKO^C* mice. Moreover, the magnitude of the protective effect of *Dll4* astrocyte knockdown was stronger in *Dll4^ACKO^C* mice than in *Dll4^ACKO^P* mice.

### Astrocyte-specific *Dll4* inactivation induces down-regulation of astrocyte reactivity under neuroinflammatory condition both *in vitro* and *in vivo*

To test the importance of astrocyte DLL4 expression at the Glia limitans during neuroinflammation, we conditionally disrupted DLL4 expression in human astrocytes, and examined the consequences on astrocyte reactivity. To do so, we used human Normal Astrocytes (NA) from ScienCell that we transfected with either a *siRNA control* or a *siRNA* targeting *Dll4* expression. Astrocytes were then activated using the pro-inflammatory cytokine Il-1β to induce astrogliosis *in vitro*.

We first verified the efficiency of the knockdown by measuring DLL4 gene and protein expression in primary human NA cultures transfected with the *siCTRL versus siDLL4* and treated with IL-1β, and showed that DLL4 gene and protein expression were strongly downregulated in the *siDLL4* condition compared to the *siCTRL* condition (Fig 3, A-C) along with the protein expression of JAG1 and NICD (notch1 intracellular domain) which reflects the level of activation of the notch pathway (Fig 3, D-F). The down-regulation of the DLL4-NOTCH1 axis was paired with the down-regulation of identified reactive astrocyte markers notably GFAP (glial acidic fibrillary protein) (25) and cleaved CASPASE 3 (26) (Fig 3, D, G-H). We then confirmed that *Dll4* knockdown leads to disruption of astrogliosis *in vivo*, on spinal cord lysates from EAE induced *Dll4^ACKO^P* and *Dll4^ACKO^C* mice and *control* littermates.

**Fig 3:**
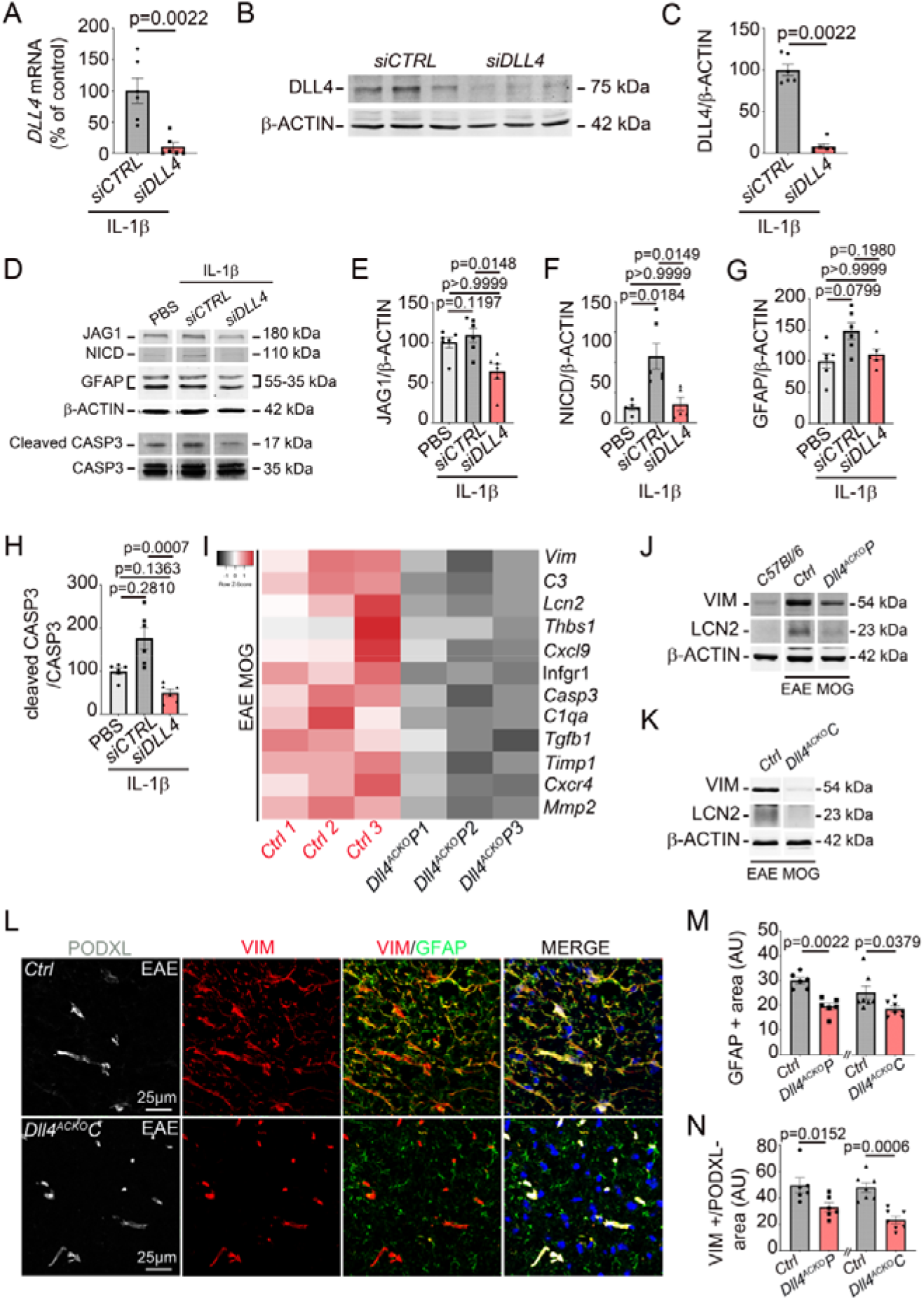
Astrocyte-specific *Dll4* inactivation induces down-regulation of astrocyte reactivity under neuroinflammatory condition both *in vitro* and *in vivo*: **(A-C)** human Normal Astrocytes (NA) were seeded and cultured until 70% confluence. They were then transfected with a *CONTROL siRNA* (20µM) *versus* a *DLL4 siRNA* (20 µM), starved for 12 hours and treated with IL-1β 10ng/mL for 12h. The experiment was repeated 3×. **(A-C)** DLL4 expression was quantified by **(A)** qRT-PCR and **(B-C)** western blot. β-ACTIN was used as a reference. **(B)** Representative western blot for DLL4 and β-ACTIN protein expression level are shown. **(C)** DLL4 protein expression level was quantified. **(D-H)** Human Normal Astrocytes (NA) were seeded and cultured until 70% confluency. They were then treated as above and **(D-E)** JAG1, **(D, F)** NICD, **(D, G)** GFAP, and **(D, H)** cleaved CASP3 expression were quantified by western blot. β-ACTIN and CASP3 total were used as references. **(I)** Spinal cord sections were harvested from MOG_35-55_ EAE induced *Dll4ACKO1* mice and *control* littermates at 18 days post induction. Transcriptional *RNA* profiling of spinal cord lysates was performed and a Heatmap generated showing the list of genes involved in astrogliosis whose expression is different in both groups. **(J-K)** VIM and LCN2 protein expression were then quantified by western blot in spinal cord lysates from 10 weeks old *C57BL/6* mice and EAE induced *Dll4ACKO1* mice, *Dll4ACKO2* mice and *control* littermates (at 18 days post induction). β-ACTIN was used as a reference. **(J-K)** Representative western blot for VIM, LCN2 and β-ACTIN in spinal cord lysates from **(J)** 10 weeks old *C57BL/6* mice and EAE induced *Dll4^ACKO^P* mice *versus controls* and **(K)** EAE induced *Dll4^ACKO^C* mice *versus controls* are shown. **(L-N)** Spinal cord EAE lesions from *Dll4^ACKO^P* mice, *Dll4^ACKO^C* mice and littermate *controls* were harvested at 18 days post induction and tissues were immune-stained with anti-GFAP (in green), anti-VIM (in red) and anti PODXL (in grey) antibodies. Nuclei were stained with DAPI (in blue). (L) *Dll4^ACKO^C* mice *versus control* tissues are shown. **(M)** GFAP+ and **(N)** VIM+/PODXL-areas were quantified (*Dll4^ACKO^P* mice *n* = 6, WT *n* = 6), **(O)** (*Dll4^ACKO^C* mice n = 7, WT n = 7)). Mann-Whitney U test or Kruskal Wallis test.

Transcriptional profiling of spinal cord lysates from EAE induced *Dll4^ACKO^P* and *control* littermates showed that 637 genes were downregulated in EAE induced *Dll4^ACKO^P* mice while 68 genes were up-regulated (Figshare DOI: 10.6084/m9.figshare.23708805). Notably, among the downregulated genes, a wide cohort of transcripts linked to glial cell activation including reactive astrocyte markers (Fig 3, I). Specifically, this approach identified, among others, *Vim* (vimentin) and *Lcn2* (lipocalin2) transcripts as downregulated in spinal cord samples from EAE induced *Dll4^ACKO^P* mice (Fig 3, I). Importantly, these markers stand today as two of the most significant reactive astrocyte markers (25) identified in the literature. We then confirmed VIM and LCN2 protein down-regulation, in spinal cord lysates from both EAE induced *Dll4^ACKO^P* and *Dll4^ACKO^C* mice compared to control littermates (Fig 3 J-K). Surprisingly, *Gfap* transcripts weren’t modulated in spinal cord samples from EAE induced *Dll4^ACKO^P* mice (Figshare DOI: 10.6084/m9.figshare.23708805). However, in examining GFAP protein expression on spinal cord sections from EAE-induced *Dll4^ACKO^P* mice and *control* littermates and from EAE-induced *Dll4^ACKO^C* mice and *control* littermates, we found that both GFAP and VIM expression are decreased in the CNS of astrocyte specific *Dll4* deficient mice (*Dll4^ACKO^* mice) (Fig 3, L-N) (Supplemental Fig 2, A).

These findings support a requirement for astrocyte DLL4 expression in the induction of astrogliosis under neuroinflammatory conditions.

### DLL4 expression depends on the phosphorylation of ERK 1/2 *in vitro* and drives NICD upregulation in astrocytes leading to the upregulation of IL-6 transcripts *via* a direct interaction with NICD

Next, we explored the signaling factors upstream and downstream of DLL4 induced astrogliosis. In the literature, Xia et al. have shown that ERK (extracellular signal-regulated kinase)-inhibition attenuated LPS (lipopolysaccharide)-induced endothelial DLL4 expression and angiogenic sprouting, both *in vitro* and *in vivo* (27). Therefore, we next measured the expression level of the phosphorylated form of P42-44 MAPK (mitogen-activated protein kinases or ERK 1/2) in reactive astrocyte cultures treated with U0126, a cell permeable inhibitor of MAPK (ERK 1/2) which blocks the kinase activity of MAP Kinase, or control vehicle. We showed that down-regulation of phospho-P42-44 (P-P42-44) is effective in U0126 treated reactive astrocytes when compared to control (Fig 4, A, D) and associated with the down-regulation of both DLL4 and NICD expression (Fig 4, A-C) suggesting that the DLL4-NOTCH1 axis activity depends on the phosphorylation of ERK 1/2 *in vitro*.

**Fig 4:**
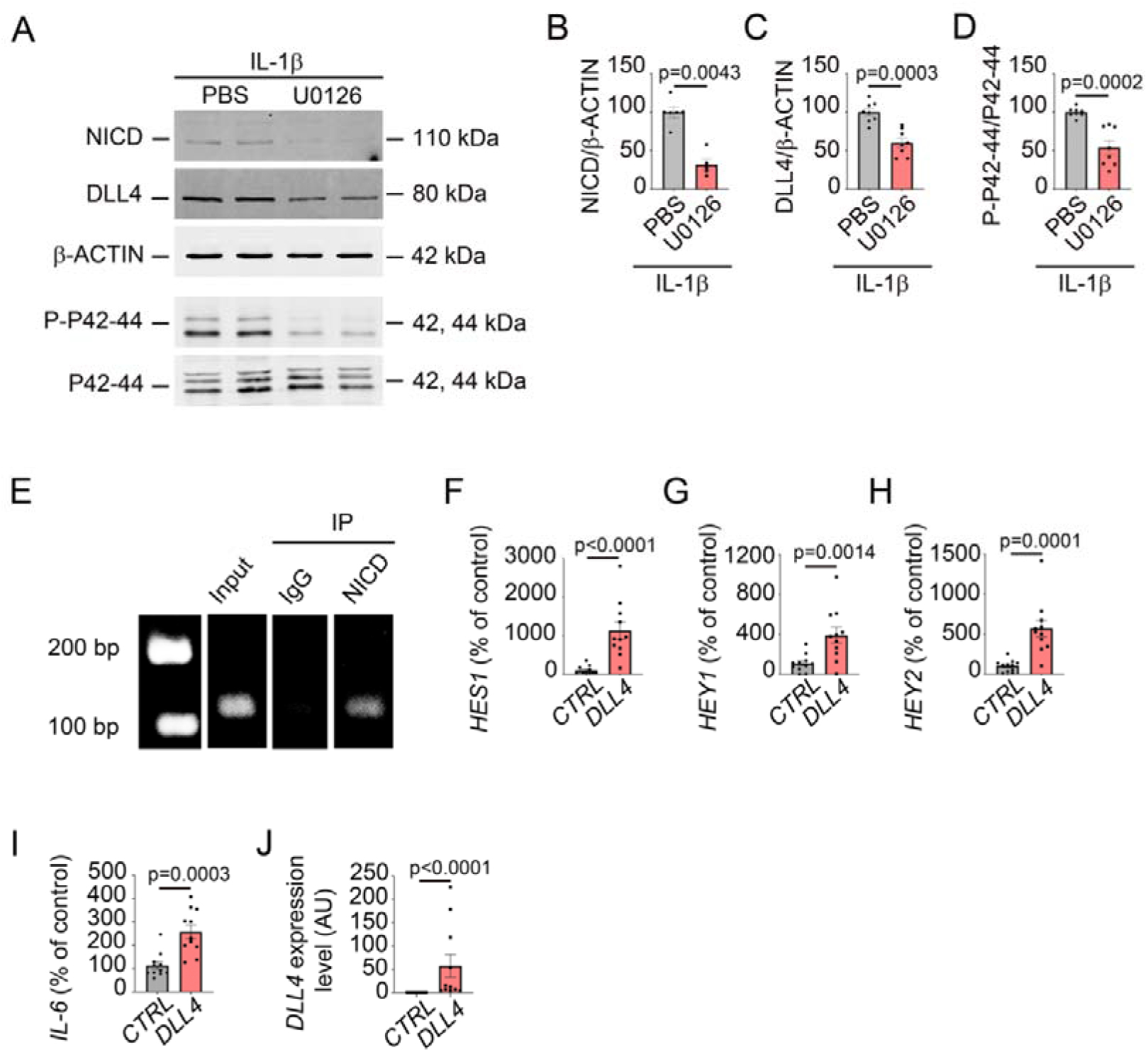
DLL4 expression depends on the phosphorylation of ERK 1/2 *in vitro* and drives NICD upregulation in astrocytes leading to the upregulation of IL-6 transcripts *via* a direct interaction with NICD: **(A-D)** Human Normal Astrocytes (NA) were seeded and cultured until confluence. They were then starved for 12 hours and treated with IL-1β 10ng/mL for 24h with or without U0126 (10 µM), a cell permeable inhibitor of MAPK (ERK ½). The experiment was repeated 3×. **(A-B)** NICD, **(A, C)** DLL4 and **(A, D)** P-P42-44 expression were quantified by western blot. β-ACTIN and P42-44 total were used as references. **(E)** Human Normal Astrocytes (NA) were seeded and cultured until confluence. They were then starved for 12 hours and treated with IL-1β 10ng/mL for 24h. A Chromatin Immuno-Precipitation (ChiP) was then performed on NA lysates using NICD antibody versus IgG controls to pull-down*. IL-6* DNA expression level was then quantified by PCR. **(F-J)** Human Normal Astrocytes (NA) were seeded and cultured until 70% confluence. They were then transduced with an empty lentivirus (6.21 10^8^ PFU/mL) *versus* a *DLL4*-expressing lentivirus (4.14 10^8^ PFU/mL) and harvested 24h post transduction. The experiment was repeated 3×. (F) *HES1*, (G) *HEY1*, (H) *HEY2*, (I*) IL-6* and (J) *DLL4* expression were quantified by qRT-PCR. β*-ACTIN* was used as a reference. Mann-Whitney U test.

Interestingly, NICD has been found to directly regulate IL-6 (interleukin-6) expression in activated macrophages (28) and our transcriptional RNA profiling of spinal cord lysates from EAE induced *Dll4^ACKO^P* and *control* littermates revealed a strong cohort of transcripts linked to IL-6 production (Figshare DOI: 10.6084/m9.figshare.23708805). Therefore we hypothesized that a direct interaction between NICD and IL-6, already observed in reactive macrophages, may contribute to reactive astrogliosis pathways *in vitro*. To respond to this question, we performed a ChIP experiment on reactive astrocyte lysates using IgG *versus* NICD antibodies to do the pull down and human *IL-6* primers for the PCR and demonstrated that NICD directly interacts with IL-6 in reactive astrocytes *in vitro* (Fig 4, E). Then, to rule out any potential effect of the inflammatory factor IL-1β on the upregulation of IL-6 in our *in vitro* model, we transduced non-reactive human astrocytes with a *DLL4* expressing lentivirus *versus* an *empty* lentivirus. First we validated that the DLL4-NOTCH1 pathway was activated in the astrocyte cultures transduced with the *DLL4* expressing lentivirus, showing the upregulation of *HES1, HEY1* and *HEY2* genes (Fig 4, F-H) and highlighted the concurrent upregulation of *IL-6* expression level (Fig 4, I). *DLL4* upregulation was also validated to ensure the transduction efficacy (Fig 4, J).

Altogether, these results suggest that DLL4 induction depends on the phosphorylation of ERK 1/2 in reactive astrocytes and that DLL4 driven NOTCH1 activation leads to the upregulation of IL-6 transcripts *via* a direct interaction between NICD and the gene coding for IL-6.

### Astrocyte-specific *Dll4* inactivation induces downregulation of astrocyte reactivity through the downregulation of the IL-6-JAK/STAT pathway both *in vitro* and *in vivo*

JAK/STAT (janus kinase/signal transducer and activator of transcription) signaling is an essential effector pathway for the development and regulation of immune responses. Unbridled activation of the JAK/STAT pathway by pro-inflammatory cytokines, notably IL-6, plays a critical role in driving the pathogenesis of multiple sclerosis/EAE (29). Moreover, STAT3 has been shown to control astrogliosis in various pathologies such as ischemic stroke, neuroinflammatory disorders and brain tumors (30–32). We tested whether the IL-6-JAK/STAT pathway was regulated by astrocytic *Dll4 in vitro* and *in vivo*. First we showed that both IL-6 and P-STAT3 (phosphorylated form of STAT3) were downregulated in reactive human astrocytes transfected with the *DLL4 siRNA* compared to reactive human astrocytes transfected with the *CTRL siRNA*. Notably, IL-6 and P-STAT3 expression level in the *DLL4 siRNA* condition was similar to the one measured in non-reactive astrocytes (Fig 5, A-C). Under the same conditions *in vitro* and on spinal cord sections from *Dll4^ACKO^C* mice and *control* littermates, we then demonstrated that IL-6 signal was co-localized with GFAP signal in reactive astrocytes (Fig 5, D-E) (Supplemental Fig 2, B) and that IL-6 expression level was downregulated following *Dll4* knockdown both *in vitro* and *in vivo* (Fig 5, D-G) (Supplemental Fig 2, B). To clearly establish the link between IL-6 and P-STAT3, we then measured the phosphorylation of STAT3 *in vitro*, in human reactive astrocytes treated with Tocilizumab, a humanized monoclonal antibody targeting IL-6 receptors, or IgG. We showed that P-STAT3 is downregulated after Tocilizumab treatment compared to IgG with an expression level similar to the one of non-reactive astrocytes (Fig 5, H-I). Importantly, DLL4 expression level was stable in both IgG and Tociliumab treated reactive astrocytes, highlighting that IL-6-JAK/STAT interaction happened downstream of DLL4 during astrogliosis (Fig 5, H, J).

**Fig 5:**
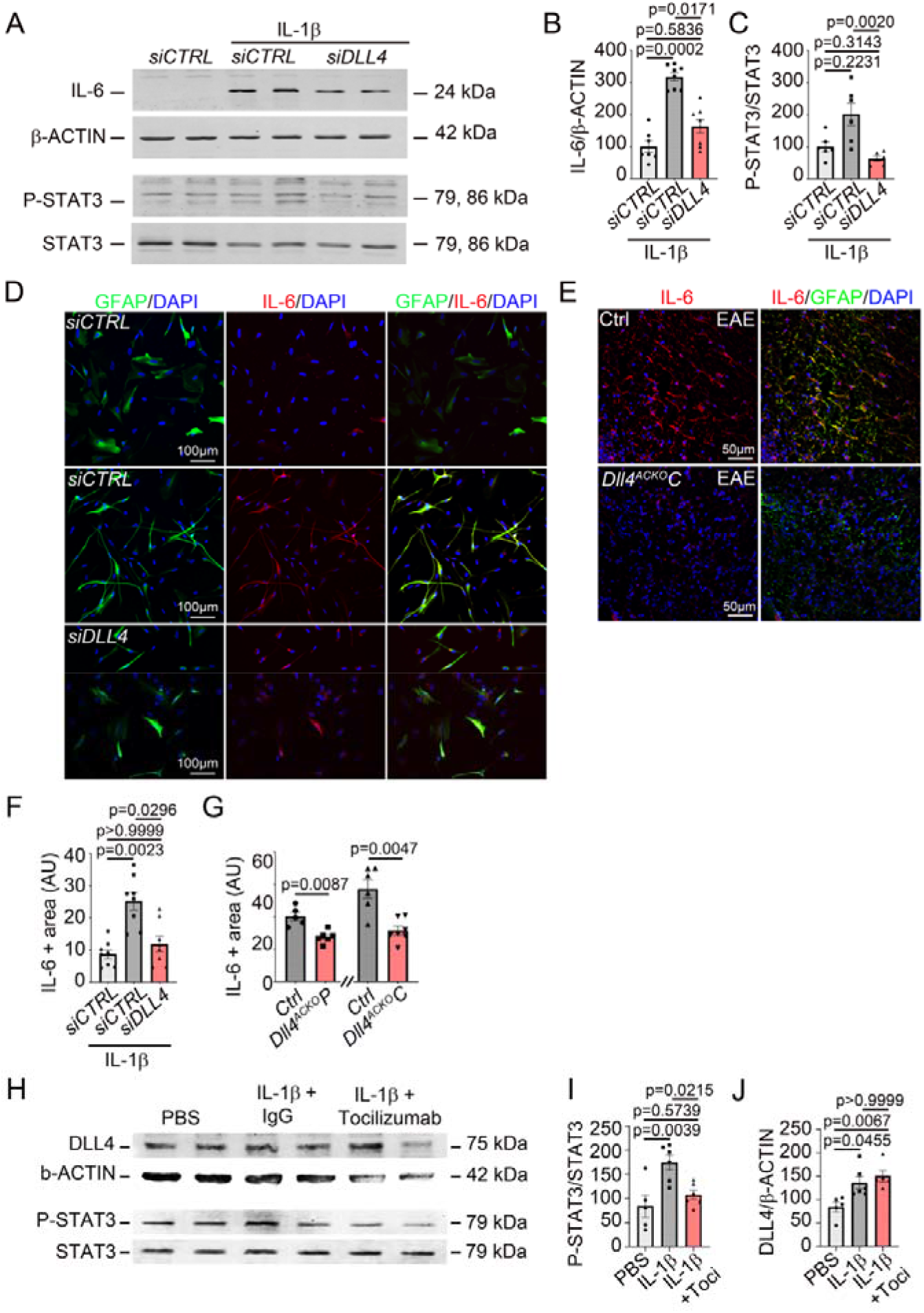
Astrocyte-specific *Dll4* inactivation induces down-regulation of astrocyte reactivity through the down-regulation of the IL-6-JAK/STAT pathway both *in vitro* and *in vivo*: **(A-C)** Human Normal Astrocytes (NA) were seeded and cultured until 70% confluence. They were then transfected with a *CONTROL siRNA* (20 µM) *versus* a *DLL4 siRNA* (20 µM), starved for 12 hours and treated with 1X PBS *versus* IL-1β 10ng/mL for 12h. The experiment was repeated 3×. **(A-B)** IL-6 and **(A, C)** P-STAT3 expression were quantified by western blot. β-ACTIN and STAT3 total were used as references**. (D)** NA were cultured on Lab-Tek® until 70% of confluence. They were then treated as above. The experiment was repeated 3×. **(D)** GFAP (in green) and IL-6 (in red) localizations were evaluated by immuno-fluorescent staining. Nuclei were stained with DAPI (in blue). **(E)** Spinal cord EAE lesions from *Dll4^ACKO^P* mice, *Dll4^ACKO^C* mice and littermate *controls* were harvested at 18 days post induction. *Dll4^ACKO^C* and *control* lesions were immune-stained with anti-GFAP (in green) and anti-IL-6 (in red) antibodies. Nuclei were stained with DAPI (in blue). *Dll4^ACKO^C* and *control* tissues are shown. **(F-G)** IL-6 positive areas were quantified **(F)** in reactive NA and **(G)** in spinal cord lesions (*Dll4^ACKO^P* mice *n* = 6, WT *n* = 5) (*Dll4^ACKO^C* mice n = 7, WT n = 6)). **(H-I)** NA were seeded and cultured until 70% confluence. They were then starved for 12 hours and treated with 1X PBS versus IL-1β 10ng/mL with IgG or an anti-IL-6 receptor antibody Tocilizumab (1µg/mL) for 24h. The experiment was repeated 3×. **(H-I)** P-STAT3 and **(H, J)** DLL4 were quantified by western blot. STAT3 total and β-ACTIN were used as references. Mann-Whitney U test OR Kruskal Wallis test

Here we showed that, following the direct upregulation of IL-6 transcription by NICD in reactive astrocytes, IL-6 protein expression is strongly increased and leads to the phosphorylation of STAT3, an already established marker of astrogliosis via its interaction with the receptor JAK.

### Mice with astrocyte *Dll4* inactivation display resistance to increases in blood brain barrier permeability via decreases in VEGFA and TYMP secretion, protecting the parenchyma from inflammatory infiltrate in a model of multiple sclerosis and in a model of acute neuroinflammation

We compared the impact of *Dll4* blockade in astrocytes on plasma protein and inflammatory cell infiltration in the parenchyma by measuring IgG, CD4 and CD45+ lymphocyte infiltration and IBA1 expression level *in vivo*, on spinal cord sections from EAE induced *Dll4^ACKO^P* and *Dll4^ACKO^C* mice versus *control* littermates (Fig 6, A-D) (Supplemental Fig 3, A-H). We found that astrocyte *Dll4* deficient mice induced with EAE displayed less parenchymal inflammatory infiltration than *control* littermates (Fig 6, A-D) (Supplemental Fig 3, A-H). We previously reported that reactive astrocytes express pro-permeability factors, VEGFA and TYMP, which drive blood brain barrier permeability in EAE (6). We therefore tested whether astrocyte DLL4 signaling regulates VEGFA and TYMP expression *in vitro* in *DLL4 siRNA* treated reactive astrocytes and *in vivo* in EAE induced *Dll4^ACKO^P* and *Dll4^ACKO^C* mice. We found that VEGFA and TYMP were downregulated in IL-1β induced reactive astrocytes transfected with *DLL4 siRNA* when compared to IL-1β induced reactive astrocytes transfected with *CTRL siRNA* (Fig 6, E-H). Moreover we showed that TYMP expression level was highly correlated to astrogliosis threshold as it followed the same pattern as GFAP signal intensity (Fig 6, H, J-K). We then confirmed these results *in vivo*, finding downregulation of astrocyte VEGFA and TYMP signals on spinal cord sections from EAE induced *Dll4^ACKO^P* and *Dll4^ACKO^C* mice when compared to *control* littermates (Fig 6, I, L-M) (Supplemental Fig 3, I-K).

**Fig 6:**
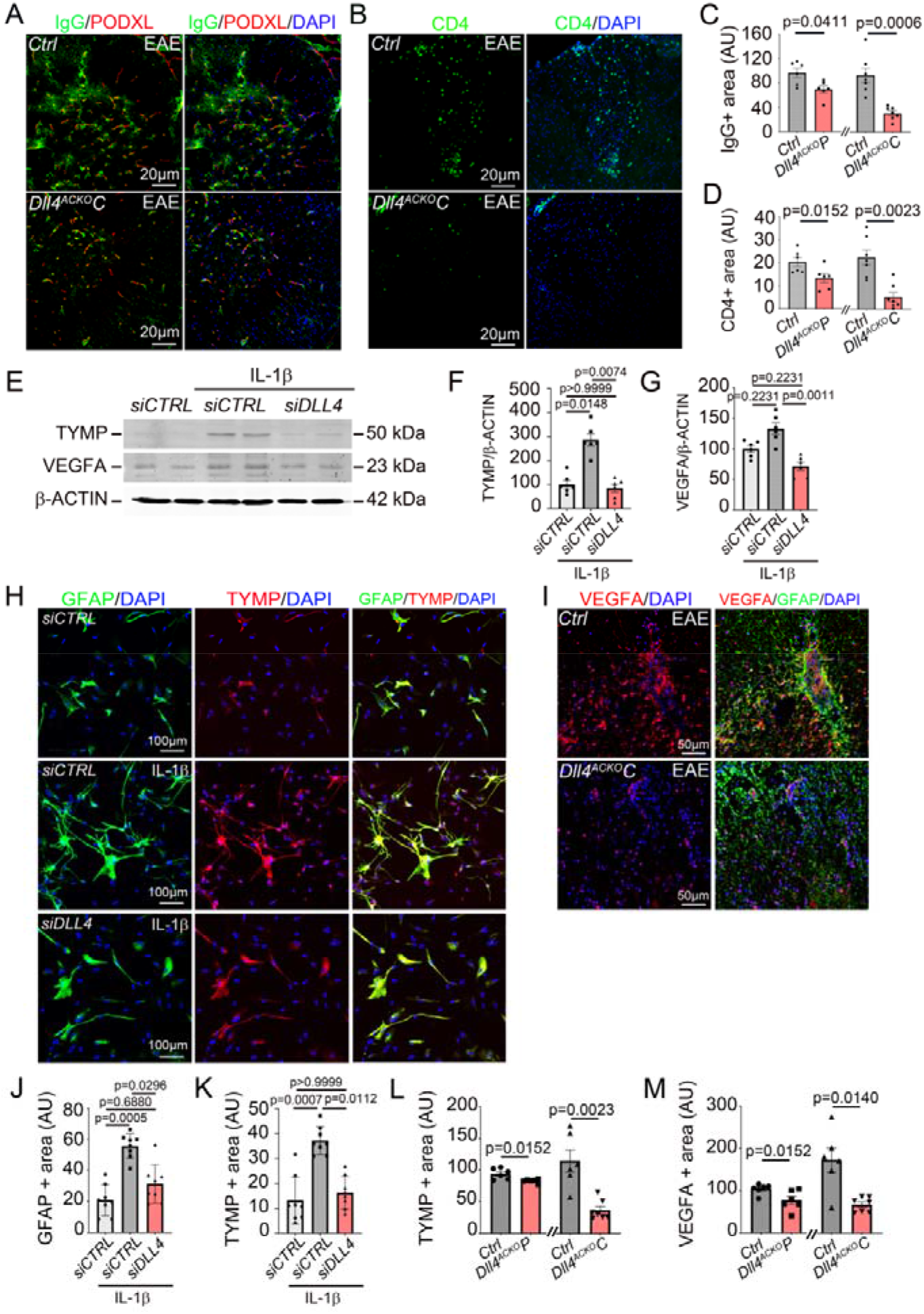
Mice with astrocyte *Dll4* inactivation display resistance to increases in blood brain barrier permeability via decreases in VEGFA and TYMP secretion, protecting the parenchyma from inflammatory infiltrate in a model of multiple sclerosis: **(A-D)** Spinal cord EAE lesions from *Dll4^ACKO^P* mice, *Dll4^ACKO^C* mice and littermate *controls* were harvested at 18 days post induction. (A-B) *Dll4^ACKO^* and *control* lesions were immune-stained with **(A)** anti IgG (in green) and anti-PODXL (in red) antibodies and **(B)** anti CD4 (in green) and anti LAM (in red) antibodies. Nuclei were stained with DAPI (in blue). *Dll4^ACKO^C* and *control* tissues are shown. **(A, C)** IgG and **(A, D)** CD4 positive areas were quantified (*Dll4^ACKO^P* mice *n* = 6, WT *n* = 5) (*Dll4^ACKO^C* mice n = 7, WT n = 7). **(E-G)** Human Normal Astrocytes (NA) were seeded and cultured until 70% confluence. They were then transfected with a *CONTROL siRNA* (20 µM) *versus* a *DLL4 siRNA* (20 µM), starved for 12 hours and treated with 1X PBS *versus* IL-1β 10ng/mL for 12h. The experiment was repeated 3×. **(E-F)** TYMP and **(E, G)** VEGFA expression were quantified by western blot. β-ACTIN was used as a reference**. (H)** Human Normal Astrocytes (NA) were cultured on Lab-Tek® until 70% of confluence. They were then treated as above. The experiment was repeated 3×. GFAP (in green) and TYMP (in red) localizations were evaluated by immuno-fluorescent staining. Nuclei were stained with DAPI (in blue). **(I)** Spinal cord EAE lesions from *Dll4^ACKO^P* mice, *Dll4^ACKO^C* mice and littermate *controls* were harvested at 18 days post induction. *Dll4^ACKO^* and *control* lesions were immune-stained with anti-GFAP (in green) and anti-VEGFA (in red) or anti-TYMP (in red) antibodies. Nuclei were stained with DAPI (in blue). Representative GFAP/VEGFA staining of *Dll4^ACKO^C* and *control* lesions is shown. **(J)** GFAP and **(K)** TYMP positive areas were quantified in reactive NA in cultures and **(L)** TYMP and **(M)** VEGFA positive areas were quantified in spinal cord lesions (*Dll4^ACKO^P* mice *n* = 6, WT *n* = 6) (*Dll4^ACKO^C* mice n = 7, WT n = 6). Mann-Whitney U test or Kruskal Wallis test

Altogether, these results suggest that, under neuroinflammatory condition, astrocyte *Dll4* knockdown leads to decreased astrogliosis and is associated with reduced levels of VEGFA and TYMP, protecting against blood brain barrier breakdown and subsequent parenchymal inflammatory infiltrate.

To further support whether astrocyte DLL4 participates to inflammatory lesion pathogenesis by controlling astrogliosis, we decided to verify the results we obtained in the EAE mouse model, in a model of acute neuroinflammatory lesion. Initially, we compared responses of *Dll4^ACKO^P* mice and littermate *controls* to cortical injection of AdIL-1β, measuring the area of neuronal cell death (NEUN loss) in lesions at 7 dpi (days post injection) (Supplemental Fig 4, A, F). We then measured astrocyte reactivity (VIM and GFAP signal intensity) and associated IL-6 and TYMP expression in lesions at 7 dpi (Supplemental Fig 4, B-C, G-J). Finally, the maximal area of inflammatory infiltration into the CNS in term of parenchymal entry of serum proteins, notably FGB (fibrinogen), and CD4^+^ T helper lymphocytes was assessed in lesions at 7 dpi (Supplemental Fig 4, D-E, K-L). Importantly, these studies demonstrated that lesion formation in *Dll4^ACKO^P* mice was strongly decreased compared to littermate *controls*. Confirming efficacy and specificity of inactivation, AdIL-1β–induced lesions in *Dll4^ACKO^P* mice showed lower levels of DLL4 (Supplemental Fig 1, K). Lesion size in *Dll4^ACKO^P* mice, as measured by neuronal cell death or loss of NEUN immunoreactivity, was much narrower than in *controls* at 7 dpi (Supplemental Fig 4, A, F). Moreover, GFAP and VIM, two markers of astrocyte reactivity were strongly downregulated in *Dll4^ACKO^P* mice compared to *controls* and were associated with a decreased expression of astrogliosis marker IL-6 and pro-permeability marker TYMP (Supplemental Fig 4, B-C, G-J). There were also large decreases in the areas of cortical FGB and CD4^+^ T helper lymphocytes infiltration seen in AdIL-1β lesions in *Dll4^ACKO^P* mice at 7 dpi (Supplemental Fig 4, D-E, K-L).

We concluded that astrocyte DLL4 upregulation during acute neuroinflammation leads to astrogliosis in the lesion area and upregulation of the pro-inflammatory cytokine IL-6 and pro-permeability factor TYMP by reactive astrocytes. Lesion size is more severe in presence of astrocyte DLL4 and associated with a stronger inflammatory infiltrate of fibrinogen and immune cells into the parenchyma. Altogether our findings in both cortical AdIL-1β lesions and spinal cord EAE support a global role for astrocytic Dll4 in driving astrogliosis and neuroinflammatory lesion pathogenesis.

### Blockade of DLL4 protects against blood brain barrier opening and paralysis in EAE

To test the therapeutic potential of exogenous DLL4 blockade during EAE, we designed experiments to test the effects of an anti-DLL4 antibody. To probe for possible off-target effects of Dll4 blockade on the vascular endothelium, we first tested the EAE phenotype in a transgenic mouse with conditional endothelial *Dll4* inactivation as DLL4 contributes to the regulation of angiogenesis *via* DLL4-mediated NOTCH1 signaling in endothelium, a key pathway for vascular development (33). Therefore, we induced EAE in 10 weeks old *Cadherin5-Cre^ERT2^, Dll4^Flox/Flox^*mice and corresponding littermate *controls* and found that *Dll4* endothelial specific down-regulation had no impact on EAE disease severity (Supplemental Fig 3, A), associated peak (Supplemental Fig 3, B) and average score during time of disability (Supplemental Fig 3, C), suggesting that exogenous Dll4 blockade with an anti-Dll4 antibody would have minimal effects on the vascular endothelium impacting the course or neuropathology of EAE.

We then compared the impact of DLL4 blockade on disease severity in EAE (Fig 7). We sensitized 8 weeks old *C57BL/6* mice (9 per group) and, beginning on day 8 post sensitization (just at the beginning of the onset of the disease), treated them every three days for eleven days with the inVivoMab anti-mouse DLL4 antibody (500 µg/mouse/d). The inVivoMab polyclonal Armenian hamster IgG (500 µg/mouse/d) was injected to the mice in the control group. Neurological signs and pathology were evaluated from day 1 to day 25 post sensitization. Onset of the disease occurred at day 10 post-sensitization, and ascending paralysis was first observed 24 h later, with severity in controls (polyclonal Armenian hamster IgG-treated mice) increasing until day 19, when neurologic deficit reached a plateau at a mean score of 3.3, indicating severe disease with complete hindlimb paralysis (Fig 7, A). Signs in anti-mouse DLL4 antibody-treated mice were much milder, peaking at day 16 at a mean score of just 2.4, indicative of a limp tail with moderate hindlimb weakness (Fig 7, A). Comparison of peak clinical severity in each individual was different between the anti-mouse DLL4 antibody and the polyclonal Armenian hamster IgG-treated regimens (Fig 7, B). For anti-mouse DLL4 antibody, differences in disease severity were highly significant compared with polyclonal Armenian hamster IgG-treated controls (Fig 7, C). Almost all controls (89%) but only 33% of anti-mouse DLL4 antibody-treated mice displayed complete hindlimb paralysis or worse (score ≥3) (Fig 7, A-C). No mortality at all was encountered in all cohorts. These improvements in clinical disease observed with the anti-mouse DLL4 antibody were associated with reduced tissue damage, in terms of decreased demyelination (Fig 7, D, H). They were also associated with reduced blood–brain barrier breakdown, plasmatic protein and inflammatory cell infiltration (Fig 7, E-F, J-L) and astrogliosis (Fig 7, G-H, M-N). No difference was observed in term of microglial activation (Fig 7, D, K). Downregulation of astrogliosis in the anti-mouse DLL4 antibody group was associated to a decreased expression of the pro-inflammatory cytokine IL-6 and of the pro-permeability factors TYMP and VEGFA (Fig 7, G-H, O-P).

**Fig 7:**
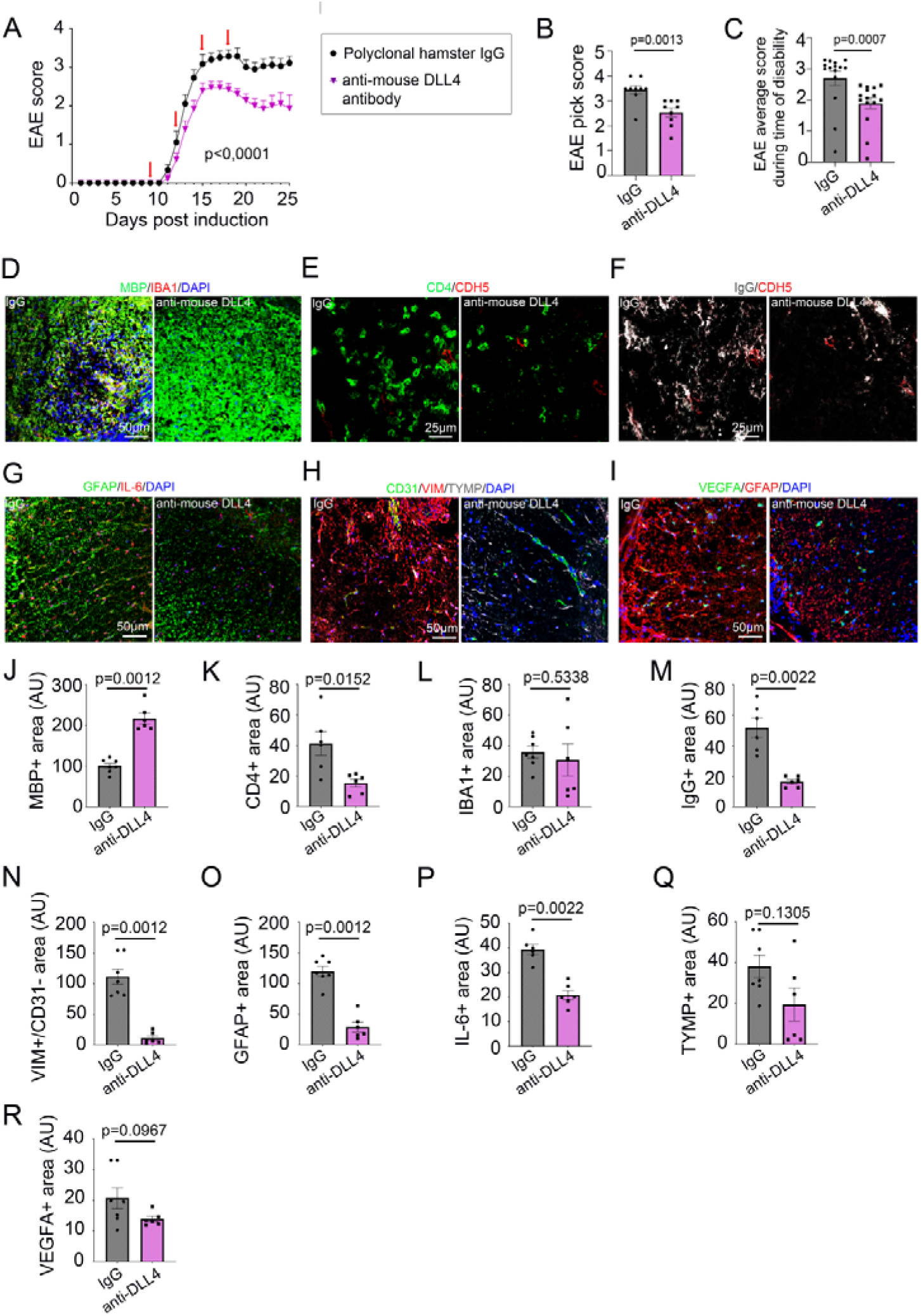
Blockade of DLL4 protects against blood brain barrier opening and paralysis in EAE: (**A**) Mean clinical scores are shown from mice (10-week-old female *C57BL/6*, 9 per group) sensitized with EAE MOG_35–55_, then from onset of disease (8 days post-induction) treated every 3 days for 11 days (red arrows) with an inVivoMab anti-mouse DLL4 antibody (500 µg/mouse/d) or an inVivoMab polyclonal Armenian hamster IgG (500 µg/mouse/d) in the control group. Neurological deficit was scored (from day 1 to day 25 post sensitization) daily for each animal according to a widely-used 5-point scale (EAE scoring: 1 limp tail; 2 limp tail and weakness of hind limb; 3 limp tail and complete paralysis of hind legs; 4 limp tail, complete hind leg and partial front leg paralysis), nonlinear regression (Boltzmann sigmoidal). **(B)** For each group of mice, EAE peak score and **(C)** EAE average score during time of disability was quantified other the course of the disease. **(D-H)** Cortical lesions were immuno-stained with **(D)** anti MBP (in green) and anti-IBA1 (in red) antibodies, **(E)** anti-CD4 (in green) and anti-CDH5 (in red) antibodies, **(F)** anti-IgG (in grey) and anti-CDH5 (in red) antibodies, **(G)** anti-GFAP (in green) and anti-IL-6 (in red) antibodies, **(H)** anti-CD31 (in green), anti-VIM (in red) and anti-TYMP (in grey) antibodies and **(I)** anti-VEGFA (in green) and anti-GFAP (in red) antibodies. **(D-I)** Nuclei were stained with DAPI (in blue). **(J)** MBP, **(K)** CD4, **(L)** IBA1**, (M)** IgG, **(N)** VIM+/CD31-, **(O)** GFAP, **(P)** IL-6, **(Q)** TYMP and **(R)** VEGFA positive areas were quantified. Mann-Whitney U test.

**Fig 8:**
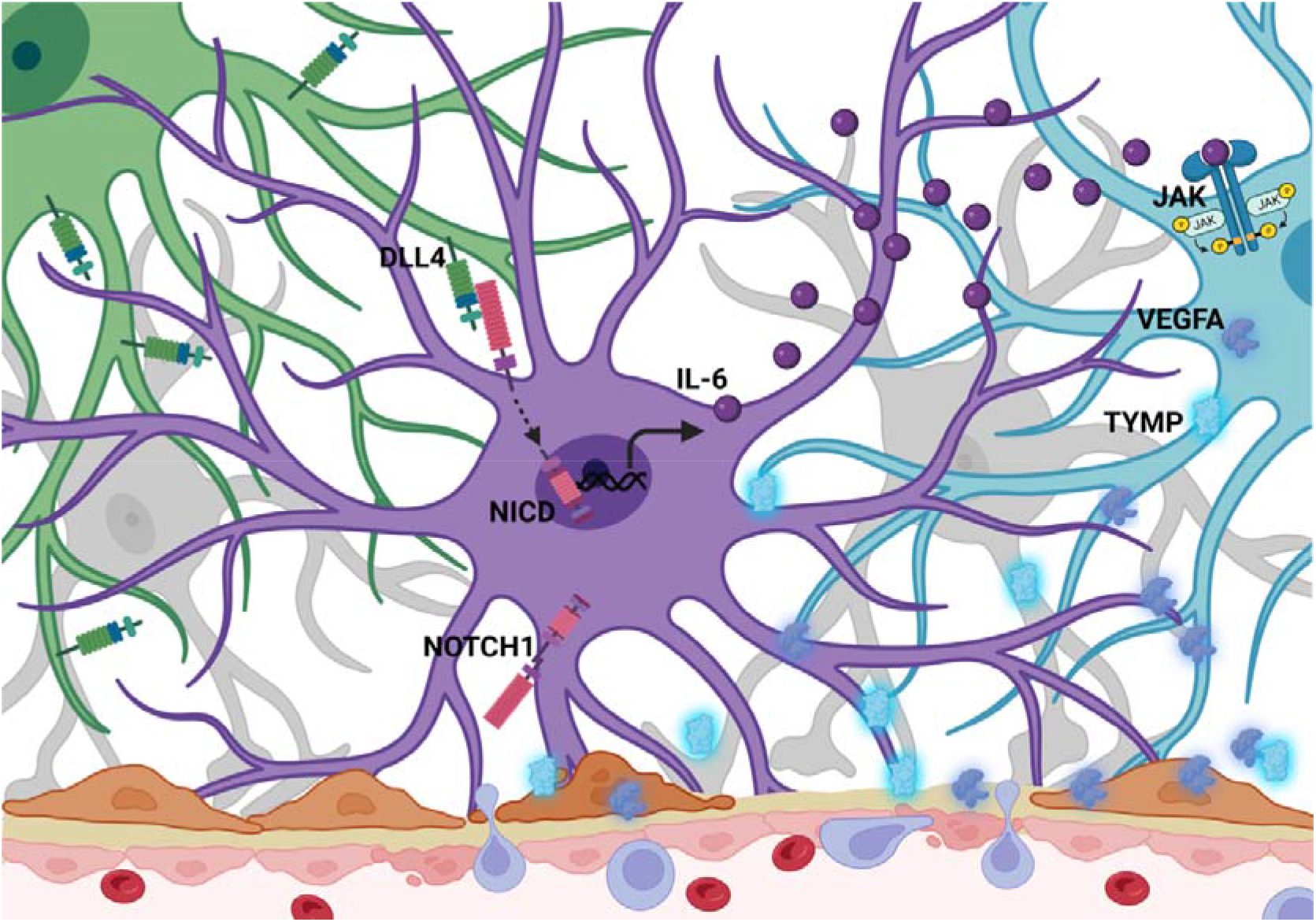
Summary scheme of the role of DLL4-NOTCH1 signaling on astrogliosis and blood brain barrier opening during neuroinflammation. In multiple sclerosis, resident and infiltrating immune cells produce cytokines that activate astrocytes. This astrogliosis leads to overexpression of DLL4 in astrocytes; DLL4 interacts with its membrane receptor NOTCH1 on the surface of a neighboring astrocytes. This leads to translocation of the intracellular active form of NOTCH1 (NICD) into the nucleus where it acts as a transcription factor to stimulate expression of the pro-inflammatory cytokine IL-6. As a result, IL-6 binds to its receptor JAK which leads to phosphorylation of STAT3, a pathway known to stimulate astrogliosis. In response to astrogliosis induced by the DLL4-NOTCH1 pathway and the NICD-IL-6-JAK/STAT signaling cascade, astrocytes secrete pro-permeability factors (TYMP and VEGFA) that participate to blood brain barrier opening and inflammatory infiltration of the parenchyma. Thus, an astrogliosis amplification loop led by DLL4-NOTCH1 is established and actively participates in the pathophysiology of neuroinflammation.

Collectively, the results of these therapies confirm the therapeutic benefit of exogenous DLL4 blockade during EAE and support our findings that astrocyte DLL4 drives astrogliosis during neuroinflammation and subsequent TYMP and VEGFA induced increases in blood brain barrier permeability.

## Discussion

A complex network of intercellular signaling occurs between cells of the neurovascular unit during neuroinflammation (1, 3). Understanding the pathways controlling reactive astrogliosis and blood brain barrier function is of considerable translational interest to the field of neuro-immunology and stroke. Here, we report for the first time, a role for juxtacrine astrocytic DLL4 mediated NOTCH1 receptor signaling as a key regulator of astrogliosis and blood brain barrier leakage in two models of CNS autoinflammatory disease.

First, we found that astrocytic DLL4 expression is up-regulated during neuroinflammation, both in mice and humans, driving astrogliosis and subsequent blood brain barrier permeability and inflammatory infiltration during pathology. We then show that the DLL4-mediated NOTCH1 juxtacrine signaling in astrocytes directly controls IL-6 transcriptional level and triggers the phosphorylation of STAT3, driving astrocyte reactivity at the Glia limitans and inducing pro-permeability factor secretion and subsequent blood brain barrier destabilization. Finally we reveal that targeting DLL4 with exogenous blocking antibodies improves EAE symptoms in mice. Together, these data identify the DLL4-NOTCH1 axis as a key driver of astrogliosis during neuroinflammation via the promotion of the IL-6-STAT3-TYMP/VEGFA signaling actors leading to the disruption of the neurovascular unit and worsening pathology.

The expression of DLL4 by reactive astrocytes has only been acknowledged once in a model of brain injury in a paper published in 2021(24). However, in the present study, we unraveled for the first time the role of the DLL4-NOTCH1 axis and associated signaling pathways, in controlling reactive astrogliosis in two models of CNS inflammation (acute CNS inflammation model and EAE, a model of multiple sclerosis). The next step would be to examine astrocytic DLL4 expression in other pathologies of the CNS, notably Alzheimer’s disease, stroke or amyotrophic lateral sclerosis in which astrocytic NOTCH1 upregulation and subsequent blood brain barrier permeability have already been identified as critical pathophysiological players (8–11). This could implicate DLL4-NOTCH1 mediated reactive astrogliosis as a more generalized mechanism in other chronic diseases of the CNS.

Like other neuroimmune factors, astrocytes are a primary source of IL-6 in the CNS (34–37) and IL-6 signaling has been shown to activate downstream pathways including JAK/STAT in various conditions including neuroinflammatory disorders^38^. In 2021, an international consortium of scientists working on astrocytes defined a “reactive astrocyte” nomenclature; they identified STAT3-mediated transcriptional programs as one of the pathways inducing astrogliosis (25). Here we demonstrated for the first time that DLL4-mediated NOTCH1 juxtacrine signaling directly controls IL-6 transcriptional level in reactive astrocytes under inflammatory condition. Based on our results and the literature, we therefore assume that DLL4-NOTCH1 axis drives reactive astrogliosis by stimulating the IL-6-JAK/STAT3 signaling cascade activity in astrocytes during neuroinflammation. Therefore, DLL4/NOTCH1 axis might be considered as a novel pathway inducing astrogliosis and could be added to the “reactive astrocyte” nomenclature as a marker of astrogliosis. It also might be interesting to look at the DLL4/NOTCH1-IL6-STAT3 signaling cascade in other chronic disorders such as Alzheimer’s and Parkinson’s diseases to broaden the scope of our study.

In the vertebrates, 4 different NOTCH receptors have been identified and at the blood brain barrier, NOTCH1 and NOTCH4 are expressed by endothelial cells (17–19) while NOTCH3 is expressed by mural cells, or pericytes (20, 21). Importantly, DLL4 was previously established as a critical regulator of angiogenesis via the DLL4-mediated NOTCH signaling in endothelial cells which is a key pathway for vascular development (33). During neuropathology, two interconnected and interdependent vascular processes are observed: blood brain barrier breakdown leading to parenchymal inflammatory infiltration (38) and abnormal angiogenesis (39). Notably, in multiple sclerosis, studies indicate that angiogenesis takes place early and show either beneficial or detrimental effect on clinical recovery depending on the literature (40). We tested extensively whether DLL4 astrocytic upregulation during neuroinflammation leads to abnormal angiogenesis and we found no differences in blood vessel density or mural cell coverage in astrocytic Dll4 knockdown mice versus controls at day 18 post-induction (data not shown). Within the neurovascular unit, astrocytes at the Glia limitans are not in direct contact with endothelial cells at the blood brain barrier and therefore, any potential regulation of angiogenesis through the DLL4-NOTCH pathway during neuroinflammation would require paracrine signaling between DLL4 secreted by astrocytes and its receptor NOTCH1 at the endothelium. We performed additional experiments testing for the presence of DLL4 expressing exosomes in conditioned media from cultured IL-1β treated human astrocytes and did not detect these (data not shown). These negative results and our current neuropathological results suggest that astrocyte DLL4 upregulation drives endothelial blood brain barrier breakdown rather than angiogenesis, via the secretion of the pro-permeability factors TYMP and VEGFA (6).

The question remains as to whether astrocytic DLL4 engages with NOTCH receptors of other cell types within the neurovascular unit, including mural cells, or pericytes, microglial processes and most interestingly, immune cells. Once immune cells penetrate the blood brain barrier, they accumulate within the perivascular space, a region between the basal basement membrane of the endothelial cell wall and the parenchymal basement membrane abutting the astrocyte endfeet (41, 42). Interestingly, NOTCH signaling can influence the differentiation and function of T lymphocytes(43). Regulatory T cells (Treg) differentially express NOTCH3 and NOTCH4 (43) and the differentiation and expansion of Tregs are reportedly induced by JAGGED engagement (44) and NOTCH3 activation (45). Moreover, novel emerging properties of the DLL4-NOTCH signaling pathway have recently been identified in pathology, notably its role in controlling the CD4+/CD8+, Th17/Treg balance both in experimental autoimmune uveitis and EAE models (46, 47).

Comparing conditional *DLL4* knockout mice and controls, we found that astrocytic DLL4 promotes the entry of T cells into the CNS parenchyma in two *in vivo* models of CNS inflammation (EAE and IL-1β cortical stereotactic injection) and that DLL4 astrocytic deletion reduces clinical disability and histopathological damage during EAE. Collectively, these results suggest that astrocytic DLL4 does not only control astrogliosis but may also influence T cell trafficking behaviors and/or activation and differentiation patterns. The extent to which T-cell activation and differentiation is influenced by local DLL4-NOTCH signaling interactions within the perivascular space has yet to be determined and is currently under investigation by our group.

Exogenous administration of an anti-mouse DLL4 antibody improved the disease in wild type mice induced with EAE, but this amelioration was moderate and similar to the one observed in EAE induced *Dll4^ACKO^P* mice which have a moderate recombination in spinal cord astrocytes. The impact of the anti-mouse DLL4 antibody therapy on EAE pathology was less significant than the one observed in the *Dll4^ACKO^C* mice which are characterized by a full recombination efficacy in spinal cord astrocytes. This discrepancy on EAE disease severity observed between a complete astrocyte *Dll4* knockdown and the anti-mouse DLL4 antibody therapy may be due to several possible factors. First, the anti-mouse DLL4 antibody is injected systemically, possibly affecting DLL4 expression in other cell types with potential opposite effects on EAE pathology. While we induced EAE in *Cadherin5-Cre^ERT2^, Dll4^Flox/Flox^* mice and littermate *controls* and showed that *Dll4* endothelial specific down-regulation has no impact on EAE disease severity, the potential still exists for effects on other unidentified cell types in adult mice induced with EAE that might express DLL4 and have opposing effects on EAE. Second, another possibility could be that the blood brain barrier, although permeable during EAE, still displays barrier properties limiting the amount of anti-mouse DLL4 antibody accessing the perivascular space and parenchyma. A third consideration is the specificity of the *Aldh1L1-Cre* promoter to astrocytes. While prior work showed high specificity of this promoter within the CNS to astrocytes, it is possible that it induces recombination in other cell types in the body expressing Dll4 which might contribute to the EAE phenotype (48).

Overall, our study identifies the DLL4-NOTCH1 axis as a key driver of astrogliosis during neuroinflammation via the IL-6-STAT3-TYMP/VEGFA signaling pathway, which is associated with disruption of the neurovascular unit and worsening pathology. This work also raises exciting questions about the role of the DLL4-NOTCH signaling in T-cell activation and differentiation (CD4+/CD8+, Th17/Treg balance) within the perivascular spaces of the neurovascular unit and its impact on neuropathology during neuroinflammation. In conclusion, this work highlights a novel role for the astrocyte ligand DLL4 in the driving of astrogliosis and breakdown of the blood brain barrier during neuroinflammation via juxtacrine NOTCH1 mediated signaling to neighboring astrocytes.

## Methods

### Human tissues

Cortical sections from multiple sclerosis patients (active lesions) and healthy controls (frontal cortex) were obtained from the Neuro-CEB bio bank (https://www.neuroceb.org/fr). The sections were 30 µm thick and obtained from fresh frozen samples.

### Mice

The *Glast-Cre^ERT2^* mice, *Aldh1L1-Cre^ERT2^* mice, *Cdh5-Cre^ERT2^* mice, *Dll4 Floxed (Dll4^Flox^)* mice and *C57BL/6* mice were purchased from Jackson Laboratories (Bar Harbor, ME, USA). The Cre recombinase in *Glast-Cre^ERT2^* mice, *Aldh1L1-Cre^ERT2^* mice *and Cdh5-Cre^ERT2^*mice was activated by intraperitoneal injection of 1 mg tamoxifen (Sigma Aldrich, St. Louis, MO, USA) for 5 consecutive days at 8 weeks of age. Mice were phenotyped 2 weeks later. Successful and specific activation of the Cre recombinase has been verified by measuring recombination efficacy in *Glast-Cre^ERT2^;Rosa26^mTmG^*mice, *Aldh1L1-Cre^ERT2^;Rosa26^mTmG^*, and *Cdh5-Cre^ERT2^; Rosa26^mTmG^* mice (Supplemental Fig 1, A-H)(4).

### Neurovascular fraction enrichment from mouse CNS

Mouse was sacrificed by cervical dislocation. Spinal cord was then harvested and transferred in a potter containing 2 mL of buffer A (HBSS 1X w/o phenol red (Gibco, Waltham, MA, USA), 10 mM HEPES (Gibco, Waltham, MA, USA) and 0,1 % BSA (Sigma Aldrich, St. Louis, MO, USA) and the tissue was pounded to obtain an homogenate which was collected in a 15 mL tube. The potter was rinsed with 1 mL of buffer A which was added to the 2 mL homogenate. Cold 30 % dextran solution was then added to the tube (V:V) to obtain a 15 % dextran working solution centrifuged for 25 minutes at 3000 g, 4 °C without brakes. After centrifugation, the pellet (neurovascular components and red cells) was collected and the supernatant (dextran solution and neural components) was centrifuged again to get the residual vessels. Neurovascular components were then pooled and re-suspended in 4 mL of buffer B (HBSS 1X Ca^2+^/ Mg^2+^ free with phenol red (Gibco, Waltham, MA, USA), 10 mM HEPES (Gibco, Waltham, MA, USA) and 0,1 % BSA (Sigma Aldrich, St. Louis, MO, USA)).

#### Neurovascular fraction enrichment for qRT-PCR

After centrifugation of the cell suspension, the pellet was washed 3 times with the buffer B and filtered through a 100 µm nylon mesh (Millipore Corporation, Burlington, MA, USA). The nylon mesh was washed with 7 mL of buffer B to collect the retained enriched neurovascular fractions. The suspension was then centrifuged for 10 minutes at 1000 g and the pellet suspended in 300 µL of RIPA lysis buffer for western blot analysis or 1000 µL of Tri-Reagent (MRC, Cincinnati, OH, USA) for qRT-PCR analysis.

### RNA sequencing

RNA was isolated using Tri Reagent® as instructed by the manufacturer (Molecular Research Center Inc). Each sample was obtained from a single mouse. mRNA library preparation was performed based on manufacturer’s recommendations (KAPA mRNA HyperPrep Kit [ROCHE]). Pooled library preparations were sequenced on NextSeq® 500 whole genome sequencing (Illumina®), corresponding to 2×30million reads per sample after demultiplexing. Quality of raw data was evaluated with FastQC(49). Poor-quality sequences were trimmed or removed with Trimmomatic(50) software to retain only good quality paired reads. Star v2.5.3a(51) was used to align reads on mm 10 reference genome using standard options. Quantification of gene and isoform abundances was done using RNA-Seq by Expectation-Maximization RSEM) 1.2.28, prior to normalization using the edgeR Bioconductor software package^39^. Finally, differential analysis was conducted with the generalized linear model (GLM) framework likelihood ratio test from edgeR. Multiple hypothesis-adjusted p-values were calculated using the Benjamini-Hochberg procedure to control for the false discovery rate (FDR). The fragments per kilobase per million mapped reads (FPKM) values of all transcripts of which expression was significantly different between MOG_35-55_ EAE-sensitized *Dll4^ACKO^P* mice and littermate *controls* is provided in Figshare (DOI: 10.6084/m9.figshare.23708805)

### Cell culture

Human Normal Astrocytes (NA) (ScienCell, Carlsbad, California) were cultured in astrocyte medium (AM) (ScienCell, Carlsbad, California). Cell from passage 2 to passage 4 were used. Before any treatment, cells were serum starved in DMEM 1 g/L glucose, Mg^+^, Ca^2+^ (Gibco, Waltham, MA, USA) without serum for 12h or 24 hours depending on the experiment.

### Transfection/Transduction

*ON-TARGET plus SMART Pool Human non-targeting* (20 µM) and *Human DLL4 siRNA* (20 µM) were purchased from Dharmacon (Colorado, USA). Cells were transfected using JetPRIME™ transfection reagent (Polyplus Transfection) according to the manufacturer’s instructions. More precisely, 500 000 astrocytes were seeded in 3.8 cm² cell culture wells. Transfection was done at day +1 with 1.8 µL of each siRNA and 4 μL JetPRIME per well. Cells were starved in DMEM 1 g/L glucose, Mg^+^, Ca^2+^ (Gibco, Waltham, MA, USA) at day +2, treated with PBS or IL-1β (10 ng/mL) at day +3 and RNA or protein samples harvested at day +4.

The human *DLL4* plasmid vector was obtained from the Theodor-Boveri-Institute, (Physiological Chemistry I, Biocenter of the University of Wurzburg) and the lentiviral plasmid was then produced and amplified in our lab. The empty lentiviral vector (*pRRLsin.MND.MCS.WPRE* (*lentiMND14*)) was produced and amplified in our lab. 500 000 astrocytes were seeded in 3.8 cm² cell culture wells and incubated for 18h before infection. Lentiviral supernatant was added with multiplicity of infection (MOI) at 10. After 8h, the lentiviral supernatant was replaced with fresh complete cell culture medium and the cells were incubated for 24 h before harvesting *RNA* samples.

### Cytokines/Growth Factors/Chemicals

Human IL-1β was purchased from PeproTech (Rocky Hills, NJ, USA) and used at 10 ng/mL. U0126, a cell permeable inhibitor of MAPK (ERK 1/2) was purchased from Cell Signaling (Danvers, MA, USA) and used at 10µM. The humanized anti-IL-6 receptor antibody Tocilizumab was used at 1µg/mL.

### Antibodies

Anti-CDH5 (Goat), anti-DLL4 (goat), anti-IL-6 (goat), anti-PODXL (goat), anti-SOX9 (goat) and anti-TYMP (goat) were from R&D systems (Minneapolis, MN, USA). Anti-FGB (rabbit) was from Dako (Carpinteria, CA, USA). Anti-human-LCN2 (mouse), anti-MBP (rat) and anti-VEGFA (Rabbit) were from Abcam (Cambridge, MA, USA). Anti-CD4 (rat) and anti-GFAP (rat and rabbit) were from Thermo Fisher Scientific (Waltham, MA, USA). Anti-RNA binding fox-1 homolog 3 (NEUN) (rabbit) was from Millipore (Billerica, MA, USA). Anti-human-DLL4 (Rabbit) and anti-LAM (rabbit) were from Sigma Aldrich (St. Louis, MO, USA). Anti β-ACTIN (rabbit), anti-CASP3 (rabbit) and Cleaved CASP3 (rabbit), anti-cleaved NOTCH1 (NICD) (rabbit), anti-DLL4 (rabbit), anti-human-JAG1 (rabbit), anti-P42-44 MAPK (rabbit) and phospho-P42-44 MAPK (rabbit), and anti-VIM (rabbit) were from cell signaling (Danvers, MA, USA). Anti-human-IL-6 (rabbit) was from Proteintech (Rosemont, IL, USA). Anti-human-STAT3 (mouse) and anti-human-P-STAT3 Tyr705 (rabbit) were from Ozyme (Saint-Cyr-l’École, France). Anti-IBA1 (rabbit) was from Fujifilm Wako (Osaka, Japan). Anti-GFP (Goat) was from Novus Biologicals (Minneapolis, MN, USA).

### Chromatin ImmunoPrecipitation

Chromatin immune-precipitation (ChIP) assay was performed with human NAs treated with IL-1β for 12h. Proteins were crosslinked to DNA by adding formaldehyde to the culture media to a final concentration of 1% and incubating for 10 minutes at 37°C. Fixed cells were rinsed twice with ice cold PBD containing AEBSF 1/1000 v/v, leupeptine 1/1000 v/v and aprotinine 1/1000 v/v. Cells (1 million) were lysed in 1.000µL of SDS buffer (SDS 1% v/v, EDTA 10mM, Tris-HCl 50mM pH8, AEBSF 1/1000 v/v, leupeptine 1/1000 v/v, aprotinine 1/1000 v/v). Lysate was sonicated on ice 20 times for 10 seconds to obtain DNA fragments with an average size of 500bp. The chromatin solution was diluted in dilution buffer to a final volume of 2mL and incubated 30 minutes at +4 °C with 30µg of protein G sepharose and 5µg of salmon sperm DNA. After centrifugation, the supernatant was incubated overnight at +4°C with 10µl of anti-human NICD antibody or non-specific IgG. After being washed twice in PBS, 30µg of G protein beads were blocked with 5µg of salmon sperm DNA added to each sample and incubated one hour at +4°C. Immuno-precipitated samples were centrifuged and washed once in low salt wash buffer, once in high salt wash buffer, once in LiCl wash buffer. Samples were then washed twice in Tris-EDTA. Elution of immune-precipitated chromatin was performed by adding 250µL of elution buffer and incubation was performed at room temperature for 15 minutes, and repeated once. To reverse DNA crosslinking, the eluate was incubated for 4 hours at 65°C after adding 20µL of NaCl 5M. Samples were then incubated 1 hour at 65°C with 2µL of proteinase K, 10µL of EDTA 0,5M and 20µL of TE 1M pH 6.5 for protein removal. DNA was then purified using phenol/chloroform extraction.

### Quantitative RT-PCR

RNA was isolated using Tri Reagent® (Molecular Research Center Inc) as instructed by the manufacturer, from 3×10^5^ cells or from isolated mouse enriched neurovascular fractions. For quantitative RT-PCR analyses, total RNA was reverse transcribed with M-MLV reverse transcriptase (Promega, Madison, WI, USA) and amplification was performed on a DNA Engine Opticon®2 (MJ Research Inc, St Bruno, Canada) using B-R SYBER® Green SuperMix (Quanta Biosciences, Beverly, MA, USA). Primer sequences are reported in Table 1.

**Table 1:**
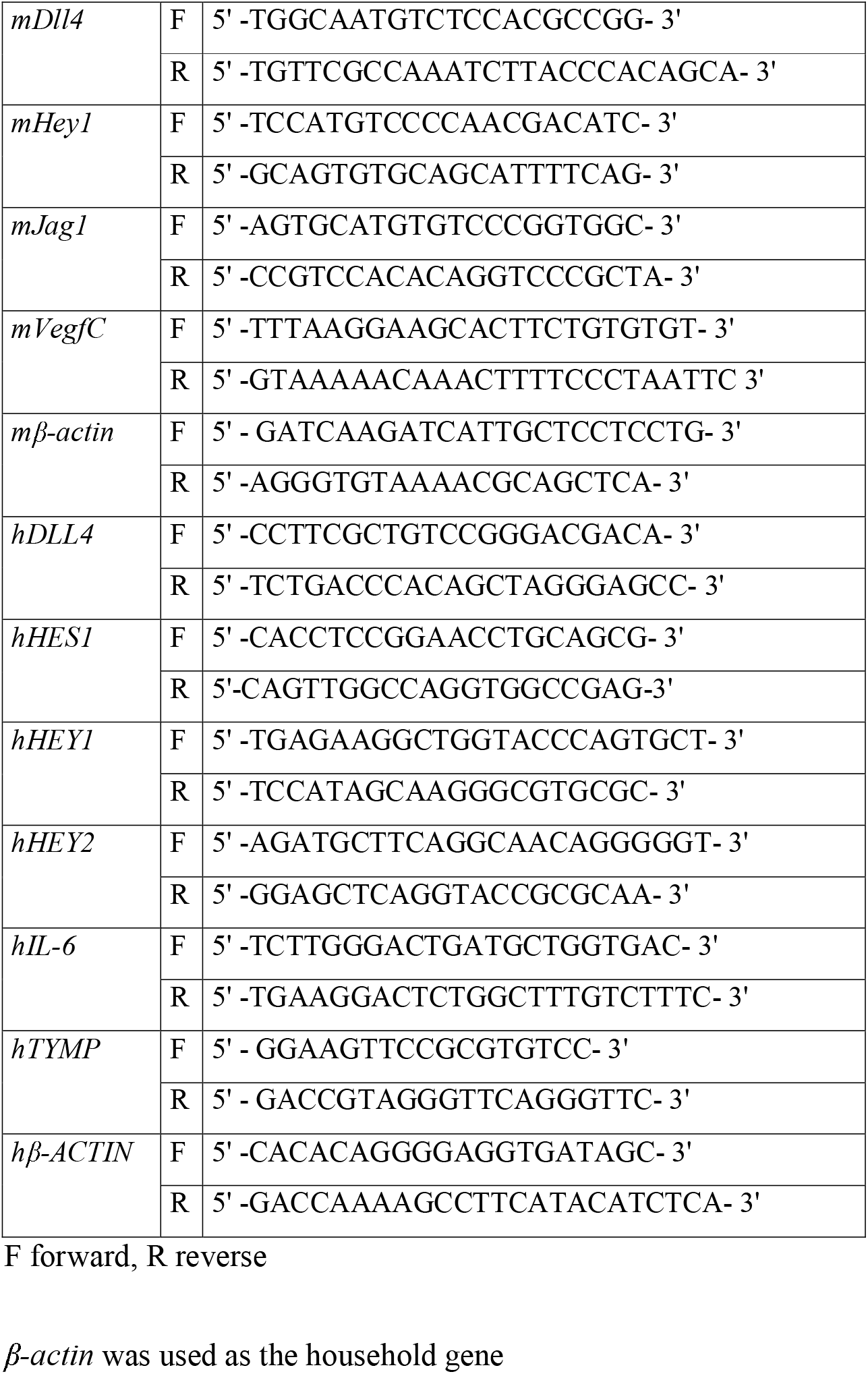
List of primers used for reverse transcription (RT) quantitative polymer chain reaction (qPCR)

The relative expression of each mRNA was calculated by the comparative threshold cycle method and normalized to β*-actin* mRNA expression.

### Western Blots

Protein expression was evaluated by SDS-PAGE. Protein loading quantity was controlled using the rabbit monoclonal anti-β-actin antibody (cell signaling, Danvers, MA, USA). Secondary antibodies were from Invitrogen. The signal was then revealed by using an Odyssey Infrared imager (LI-COR, Lincoln, NE, USA). For quantification, the mean pixel density of each band was measured using Image J software (NIH, Bethesda, MD, USA) and data were standardized to β-actin, and fold change versus control calculated.

### Stereotactic injection

10 weeks old *Dll4^ACKO^P* mice and littermate *controls* (6 mice per condition) were anaesthetized using isoflurane (3% induction and 1% maintenance) (Virbac Schweiz, Glattbrugg, Germany) and placed into a stereotactic frame (Stoelting Co., IL, USA). To prevent eye dryness, an ophthalmic *ointment* was applied at the ocular surface to maintain eye hydration during the time of surgery. The skull was shaved and the skin incised on 1 cm to expose the skull cap. Then, a hole was drilled into the skull, using a pneumatic station S001+TD783 Bien Air, until reaching the dura mater. AdIL-1 (10^7^ PFU) was then delivered into the frontal cortex at coordinates of 1 µm posterior to bregma, 2 µm left of the midline and 1.5 µm below the surface of the cortex.

Mice received a subcutaneous injection of buprenorphine (0,05 mg/kg) (Ceva santé animale, Libourne, France) 30 minutes before surgery and again 8 hours post-surgery to assure a constant analgesia during the procedure and postoperatively. Mice were sacrificed by pentobarbital (Richter Pharma, Wels, Austria) overdose at 7 days post injection (dpi). For histological assessment, the brain of each animal was harvested.

### Experimental autoimmune encephalomyelitis (EAE)

10 week old female mice were immunized by subcutaneous injection of 300 µg myelin oligodendrocyte glycoprotein-35-55 (MOG_35–55_) (Hooke laboratories, Lawrence, MA, USA) in 200 µl Freund’s Adjuvant containing 300 µg/mL mycobacterium tuberculosis H37Ra (Hooke laboratories, Lawrence, MA, USA) in the dorsum. Mice were administered with 500 ng pertussis toxin (PTX) *intra-peritoneously* on day of sensitization and 1 day later (Hooke laboratories). The emulsion provides antigen which initiates expansion and differentiation of MOG-specific autoimmune T cells. PTX enhances EAE development by providing additional adjuvant. EAE will develop in mice 7-14 days after immunization (Day 0): animals which develop EAE will become paralyzed. Disease was scored (0, no symptoms; 1, floppy tail; 2, hind limb weakness (paraparesis); 3, hind limb paralysis (paraplegia); 4, fore- and hind limb paralysis; 5, death)(52) from day 7 post immunization until day 28 post immunization. At Day 28, all the animals were euthanized by pentobarbital (Richter Pharma, Wels, Austria) overdose. For histological assessment, cervical, lumbar and dorsal sections of each animal spinal cord, as well as the spleen, were harvested.

### Therapies

inVivoMab polyclonal Armenian hamster IgG (500 µg/mouse/d) was used as control group therapy. InVivoMab anti-mouse DLL4 antibody (500 µg/mouse/d) was used as experimental group therapy. Intra-peritoneal injection of each therapy was injected (9 mice per therapy) at day 8, day 12, day 15 and day 18 post EAE immunization.

### Immunohistochemistry

Prior to tissue collection and staining, mice were transcardially perfused with PBS (10mL) followed by 10% Formalin (10mL) to remove intravascular plasma proteins. Brain and spinal cord samples were either fixed in 10 % formalin for 3 hours, incubated in 30 % sucrose overnight, OCT embedded and cut into 9 µm thick sections or directly OCT embedded and cut into 9 µm thick sections. Cultured cells were fixed with 10 % formalin for 10 minutes. Human frozen sections were used directly without any prior treatment. Concerning the fixed sections, for IL-6, prior to blocking, sections were soaked in Citrate (pH 7.5; 100 °C). For TYMP, prior to blocking, sections were soaked in EDTA (pH 6.0; 100 °C). For CD4, sections were treated with 0.5 mg/mL protease XIV (Sigma Aldrich, St. Louis, MO, USA) at 37 °C for 5 minutes. Primary antibodies were used at 1:100 except DLL4 (1:50), FGB (1:500) and LAM (1:1,000). Samples were examined using a Zeiss Microsystems confocal microscope (Oberkochen, Germany), and stacks were collected with *z* of 1 μm.

For immunofluorescence analyzes, primary antibodies were resolved with Alexa Fluor®– conjugated secondary polyclonal antibodies (Invitrogen, Carlsbad, CA, USA) and nuclei were counterstained with DAPI (1:5000) (Invitrogen, Carlsbad, CA, USA). For all immunofluorescence analyses, negative controls using secondary antibodies only were done to check for antibody specificity.

### Statistical analyses

Results are reported as mean ± SEM. Comparisons between groups were analyzed for significance with the non-parametric Mann-Whitney test, the non-parametric Kruskal Wallis test followed by the Dunn’s multiple comparison test when we have more than 2 groups or a nonlinear regression test (Boltzmann sigmoidal) for the EAE scoring analysis using GraphPad Prism v8.0.2 (GraphPad Inc, San Diego, CA, USA). Differences between groups were considered significant when p≤0.05 (*: p≤0.05; **: p≤0.01; ***: p≤0.001 ****: p≤0.0001).

### Study approval

Human samples: The Neuro-CEB bio bank and the INSERM U1034 certify that all human sections utilized for this study are ethically obtained with documented, legal permission for research use (authorization number #AC-2018-3290 obtained from the Ministry of Higher Education and Research) and in the respect of the written given consent from the source person in accordance with applicable laws and the WMA Helsinki declaration of 2013. Animal experiments were performed in accordance with the guidelines from Directive 2010/63/EU of the European Parliament on the protection of animals used for scientific purposes and approved by the local Animal Care and Use Committee of the Bordeaux University CEEA50 (IACUC protocol #16901).

### Data availability

Data are available from the corresponding author upon request.

## Author contributions

P.M. and M.L. conducted experiments, acquired data and analyzed data. C.B. conducted experiments and acquired data. P. R. conducted experiments. A.-P. G., T. C. and M.-A. R. critically revised the manuscript. C.C. designed research studies, conducted experiments, acquired data, analyzed data, provided reagents, and wrote the manuscript

## Acknowledgments

We thank Dr Mary P. Heyer for her proofreading and correction of the manuscript. We thank Sylvain Grolleau, and Maxime David for their technical help. We thank Christelle Boullé for administrative assistance.

## References

1. McConnell HL, Kersch CN, Woltjer RL, Neuwelt EA. The Translational Significance of the Neurovascular Unit. J. Biol. Chem. 2017;292(3):762–770.

2. Engelhardt B, Coisne C. Fluids and barriers of the CNS establish immune privilege by confining immune surveillance to a two-walled castle moat surrounding the CNS castle. Fluids Barriers CNS 2011;8(1):4.

3. Iadecola C. The Neurovascular Unit Coming of Age: A Journey through Neurovascular Coupling in Health and Disease. Neuron 2017;96(1):17–42.

4. Mora P et al. Blood-brain barrier genetic disruption leads to protective barrier formation at the Glia Limitans. PLoS Biol 2020;18(11):e3000946.

5. Gimsa U, Mitchison NA, Brunner-Weinzierl MC. Immune privilege as an intrinsic CNS property: astrocytes protect the CNS against T-cell-mediated neuroinflammation. Mediators Inflamm. 2013;2013:320519.

6. Chapouly C et al. Astrocytic TYMP and VEGFA drive blood-brain barrier opening in inflammatory central nervous system lesions. Brain 2015;138(Pt 6):1548–1567.

7. Nitta T et al. Size-selective loosening of the blood-brain barrier in claudin-5–deficient mice. J Cell Biol 2003;161(3):653–660.

8. Zhong J-H et al. Activation of the Notch-1 signaling pathway may be involved in intracerebral hemorrhage-induced reactive astrogliosis in rats. J Neurosurg 2018;129(3):732– 739.

9. Shimada IS, Borders A, Aronshtam A, Spees JL. Proliferating reactive astrocytes are regulated by Notch-1 in the peri-infarct area after stroke. Stroke 2011;42(11):3231–3237.

10. Nonneman A et al. Astrocyte-derived Jagged-1 mitigates deleterious Notch signaling in amyotrophic lateral sclerosis. Neurobiol Dis 2018;119:26–40.

11. Cheng Y-Y et al. Reactive Astrocytes Display Pro-inflammatory Adaptability with Modulation of Notch-PI3K-AKT Signaling Pathway Under Inflammatory Stimulation. Neuroscience 2020;440:130–145.

12. Liu X et al. IL-9-triggered lncRNA Gm13568 regulates Notch1 in astrocytes through interaction with CBP/P300: contribute to the pathogenesis of experimental autoimmune encephalomyelitis. J Neuroinflammation 2021;18(1):1–15.

13. Fortini ME. Gamma-secretase-mediated proteolysis in cell-surface-receptor signalling. Nat Rev Mol Cell Biol 2002;3(9):673–684.

14. Selkoe D, Kopan R. Notch and Presenilin: regulated intramembrane proteolysis links development and degeneration. Annu Rev Neurosci 2003;26:565–597.

15. Mumm JS, Kopan R. Notch signaling: from the outside in. Dev Biol 2000;228(2):151–165.

16. Bray SJ. Notch signalling: a simple pathway becomes complex. Nat Rev Mol Cell Biol 2006;7(9):678–689.

17. Murphy PA et al. Endothelial Notch4 signaling induces hallmarks of brain arteriovenous malformations in mice. Proc Natl Acad Sci U S A 2008;105(31):10901–10906.

18. Grigorian A, Hurford R, Chao Y, Patrick C, Langford TD. Alterations in the Notch4 pathway in cerebral endothelial cells by the HIV aspartyl protease inhibitor, nelfinavir. BMC Neurosci 2008;9(1):1–18.

19. Yu L et al. Adropin preserves the blood-brain barrier through a Notch1/Hes1 pathway after intracerebral hemorrhage in mice. J Neurochem 2017;143(6):750–760.

20. Liu H, Kennard S, Lilly B. NOTCH3 expression is induced in mural cells through an autoregulatory loop that requires endothelial-expressed JAGGED1. Circ Res 2009;104(4):466–475.

21. Gaengel K, Genové G, Armulik A, Betsholtz C. Endothelial-mural cell signaling in vascular development and angiogenesis. Arterioscler Thromb Vasc Biol 2009;29(5):630–638.

22. Qian D et al. Blocking Notch signal pathway suppresses the activation of neurotoxic A1 astrocytes after spinal cord injury. Cell Cycle 2019;18(21):3010–3029.

23. Patel M et al. Nkx6.1 enhances neural stem cell activation and attenuates glial scar formation and neuroinflammation in the adult injured spinal cord. Exp Neurol 2021;345:113826.

24. Ribeiro TN, Delgado-García LM, Porcionatto MA. Notch1 and Galectin-3 Modulate Cortical Reactive Astrocyte Response After Brain Injury. Front Cell Dev Biol 2021;9:649854.

25. Escartin C et al. Reactive astrocyte nomenclature, definitions, and future directions. Nat Neurosci 2021;24(3):312–325.

26. Wagner D-C et al. Cleaved caspase-3 expression after experimental stroke exhibits different phenotypes and is predominantly non-apoptotic. Brain Res 2011;1381:237–242.

27. Xia S, Menden HL, Korfhagen TR, Kume T, Sampath V. Endothelial immune activation programmes cell-fate decisions and angiogenesis by inducing angiogenesis regulator DLL4 through TLR4-ERK-FOXC2 signalling. The Journal of Physiology 2018;596(8):1397–1417.

28. Wongchana W, Palaga T. Direct regulation of interleukin-6 expression by Notch signaling in macrophages. Cell Mol Immunol 2012;9(2):155–162.

29. Liu Y, Gibson SA, Benveniste EN, Qin H. Opportunities for Translation from the Bench: Therapeutic Intervention of the JAK/STAT Pathway in Neuroinflammatory Diseases. Crit Rev Immunol 2015;35(6):505–527.

30. Reichenbach N et al. Inhibition of Stat3-mediated astrogliosis ameliorates pathology in an Alzheimer’s disease model. EMBO Mol Med 2019;11(2):e9665.

31. Abjean L et al. Reactive astrocytes promote proteostasis in Huntington’s disease through the JAK2-STAT3 pathway. Brain 2022;awac068.

32. Ceyzériat K, Abjean L, Carrillo-de Sauvage M-A, Ben Haim L, Escartin C. The complex STATes of astrocyte reactivity: How are they controlled by the JAK-STAT3 pathway?. Neuroscience 2016;330:205–218.

33. Hellström M et al. Dll4 signalling through Notch1 regulates formation of tip cells during angiogenesis. Nature 2007;445(7129):776–780.

34. Choi SS, Lee HJ, Lim I, Satoh J, Kim SU. Human Astrocytes: Secretome Profiles of Cytokines and Chemokines. PLOS ONE 2014;9(4):e92325.

35. Dong Y, Benveniste EN. Immune function of astrocytes. Glia 2001;36(2):180–190.

36. Farina C, Aloisi F, Meinl E. Astrocytes are active players in cerebral innate immunity. Trends Immunol 2007;28(3):138–145.

37. Nakamachi T et al. IL-6 and PACAP receptor expression and localization after global brain ischemia in mice. J Mol Neurosci 2012;48(3):518–525.

38. Engelhardt B, Ransohoff RM. Capture, crawl, cross: the T cell code to breach the blood-brain barriers. Trends Immunol. 2012;33(12):579–589.

39. Girolamo F, Coppola C, Ribatti D, Trojano M. Angiogenesis in multiple sclerosis and experimental autoimmune encephalomyelitis. Acta Neuropathologica Communications 2014;2(1):84.

40. Lengfeld J, Cutforth T, Agalliu D. The role of angiogenesis in the pathology of multiple sclerosis. Vasc Cell 2014;6(1):23.

41. Engelhardt B, Coisne C. Fluids and barriers of the CNS establish immune privilege by confining immune surveillance to a two-walled castle moat surrounding the CNS castle. Fluids Barriers CNS 2011;8(1):4.

42. Engelhardt B, Ransohoff RM. Capture, crawl, cross: the T cell code to breach the blood-brain barriers. Trends Immunol. 2012;33(12):579–589.

43. Tanigaki K, Honjo T. Regulation of lymphocyte development by Notch signaling. Nat Immunol 2007;8(5):451–456.

44. Yvon ES et al. Overexpression of the Notch ligand, Jagged-1, induces alloantigen-specific human regulatory T cells. Blood 2003;102(10):3815–3821.

45. Anastasi E et al. Expression of activated Notch3 in transgenic mice enhances generation of T regulatory cells and protects against experimental autoimmune diabetes. J Immunol 2003;171(9):4504–4511.

46. Eixarch H et al. Inhibition of delta-like ligand 4 decreases Th1/Th17 response in a mouse model of multiple sclerosis. Neuroscience Letters 2013;541:161–166.

47. Yin X et al. Activation of the Notch signaling pathway disturbs the CD4+/CD8+, Th17/Treg balance in rats with experimental autoimmune uveitis. Inflamm Res 2019;68(9):761–774.

48. Nemeth D, Luqman N, Chen L, Quan N. Aldh1l1-Cre/ER T2 is expressed in unintended cell types of the salivary gland, pancreas, and spleen.. MicroPubl Biol 2023:10.17912/micropub.biology.000832.

49. de Sena Brandine G, Smith AD. Falco: high-speed FastQC emulation for quality control of sequencing data. F1000Res 2019;8:1874.

50. Bolger AM, Lohse M, Usadel B. Trimmomatic: a flexible trimmer for Illumina sequence data. Bioinformatics 2014;30(15):2114–2120.

51. Dobin A et al. STAR: ultrafast universal RNA-seq aligner. Bioinformatics 2013;29(1):15– 21.

52. Gurfein BT et al. IL-11 regulates autoimmune demyelination. J. Immunol. 2009;183(7):4229–4240.

