## Supplemental Material for "Juxtacrine DLL4-NOTCH1 signaling between astrocytes drives neuroinflammation via the IL-6-STAT3 axis"

### Supplemental data

**Supplemental Fig 1: (Related to Fig 2) *Glast-Cre$^{ERT2}$* recombinase activation is specific but moderate in spinal cord astrocytes and cortical astrocytes whereas *Aldh1L1-Cre$^{ERT2}$* recombination is specific and strong in both spinal cord and cortical astrocytes: (A-D)** spinal cord and **(E-H)** cortical sections were harvested from *GlastCre$^{ERT2}$*, *Rosa26$^{mTmG}$* mice, *Aldh1LCre1$^{ERT2}$*, *Rosa$^{mTmG}$* mice and *control* littermates. They were then immuno-stained with **(A-C)** anti-GFP (in green) and anti-GFAP (in red) antibodies or **(E-G)** anti-GFP (in green) and anti-SOX9 (in red) antibodies. Nuclei were stained with DAPI (in blue). **(D)** The ratio of GFP+ area/GFAP+ area was then quantified in spinal cord sections from *GlastCre$^{ERT2}$*, *Rosa26$^{mTmG}$* mice and *Aldh1LCre1$^{ERT2}$*, *Rosa$^{mTmG}$* mice. **(H)** The ratio of GFP+ area/number of SOX9+ cells was then quantified in cortical sections from *GlastCre$^{ERT2}$*, *Rosa26$^{mTmG}$* mice and *Aldh1LCre1$^{ERT2}$*, *Rosa$^{mTmG}$* mice

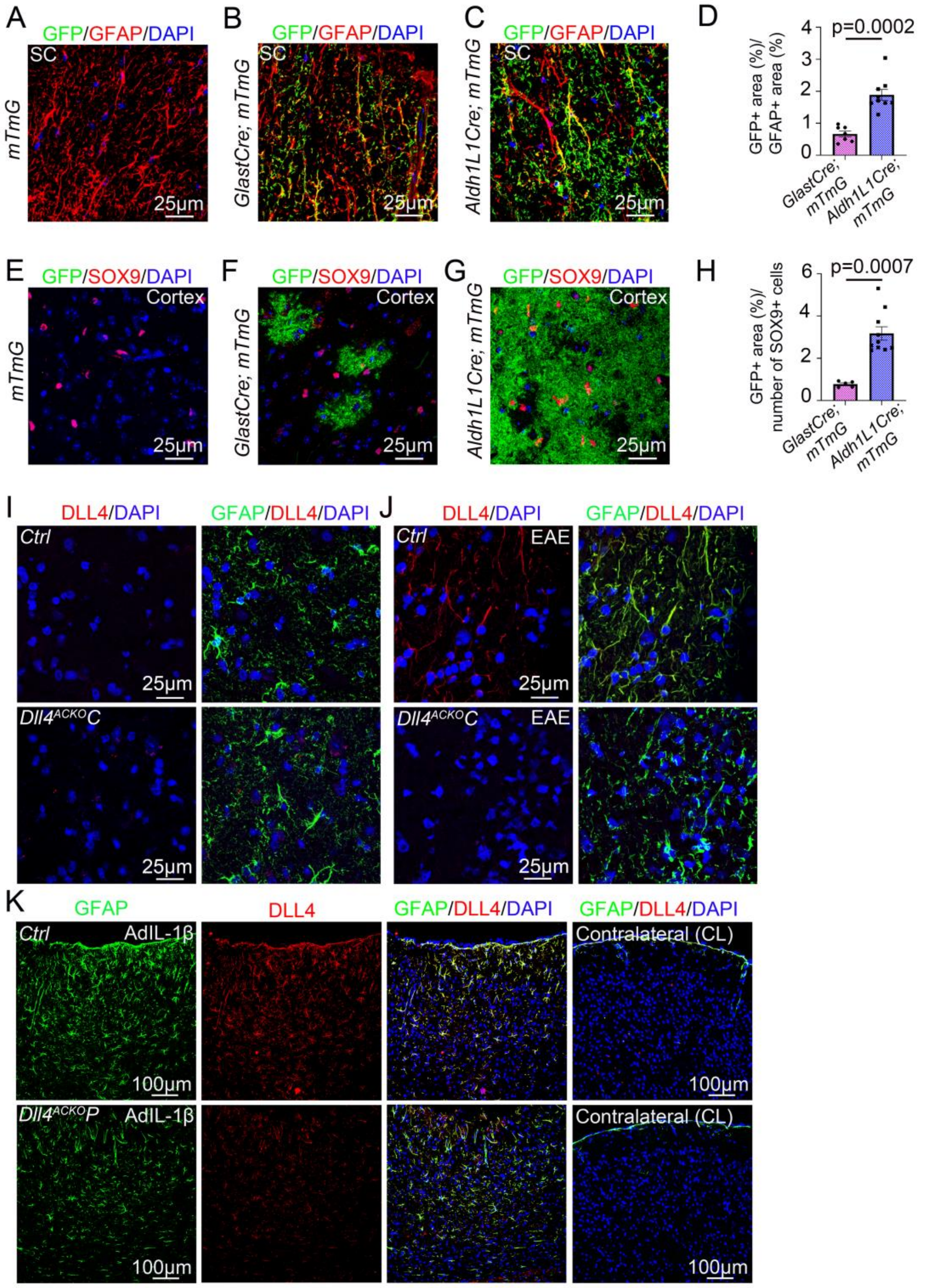


**Supplemental Fig 1: (Related to Fig 7) *Dll4* astrocyte knockdown is efficient under chronic and acute neuroinflammatory conditions: (I-J)** Spinal cord sections were harvested from **(I)** healthy versus **(J)** $MOG_{35-55}$ EAE-sensitized *Dll4$^{ACKO}$C* mice and littermate *controls* ( 18 days post induction) and immuno-stained with anti-GFAP (in green) and anti-DLL4 (in red) antibodies. **(K)** Cerebral cortices of 10-week-old *Dll4$^{ACKO}$P* mice and *control* littermates were harvested 7 days following stereotactic microinjection of AdIL-1 ($10^7$ PFU). Cortical lesions and contralateral were immuno-stained with anti-GFAP (in green) and anti-DLL4 (in red) antibodies. Mann-Whitney U test.

**Supplemental Fig 2 (Related to Fig 3):** ***Dll4$^{ACKO}$P*** **mice display less expression of GFAP and VIM under EAE condition. (A)** Spinal cord EAE lesions from *Dll4$^{ACKO}$P* mice and littermate *controls* were harvested at 18 days post induction and tissues were immune-stained with anti-GFAP (in green), anti-VIM (in red) and anti PODXL (in grey) antibodies. Nuclei were stained with DAPI (in blue). *Dll4$^{ACKO}$P versus control* lesions are shown.

**Supplemental Fig 2 (Related to Fig 5):** ***Dll4$^{ACKO}$P*** **mice display less expression of IL-6 under EAE condition. (B)** Spinal cord EAE lesions from *Dll4$^{ACKO}$P* mice and littermate *controls* were harvested at 18 days post induction. *Dll4$^{ACKO}$P* and *control* lesions were immune-stained with anti-GFAP (in green) and anti-IL-6 (in red) antibodies. Nuclei were stained with DAPI (in blue).

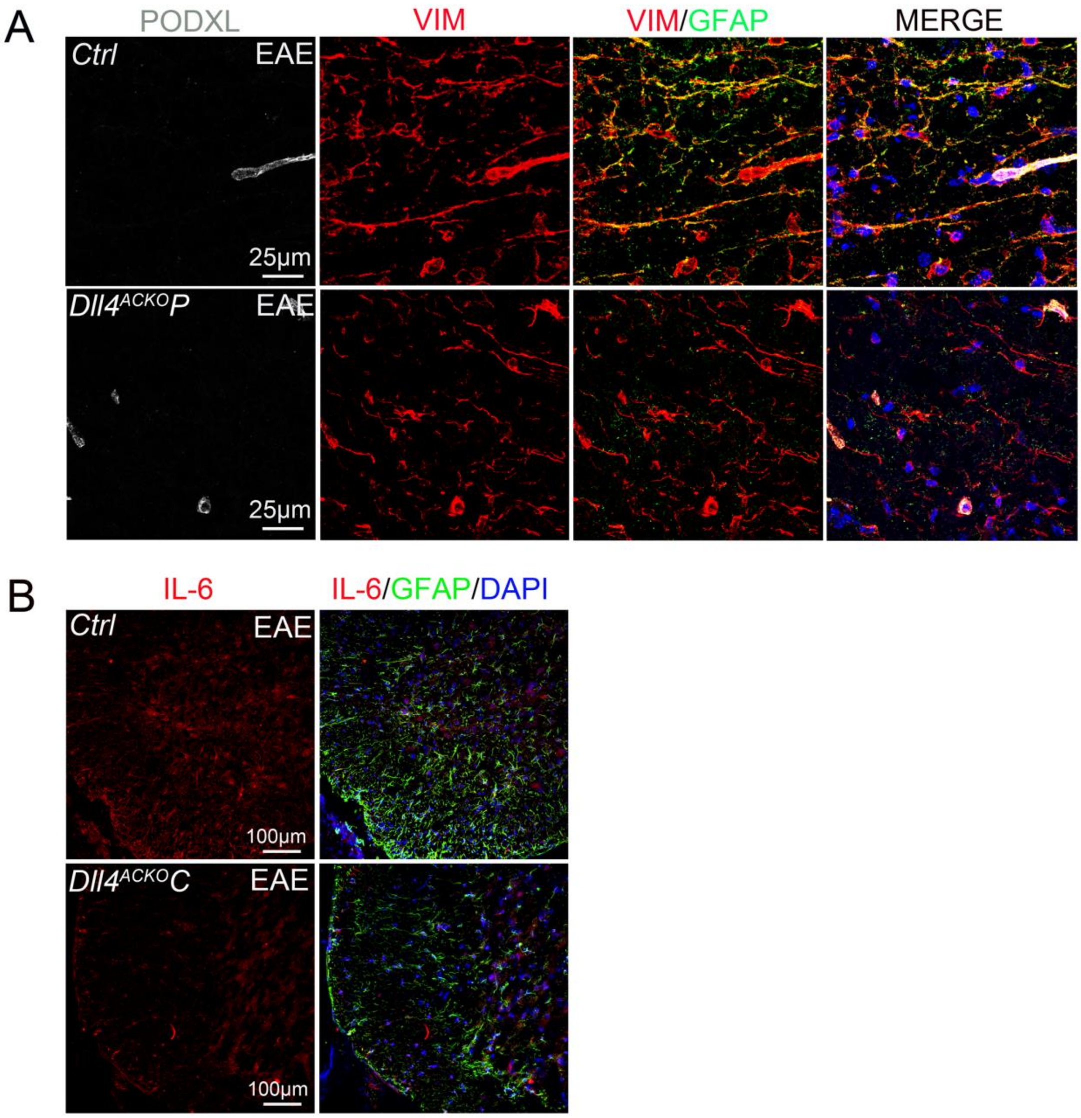

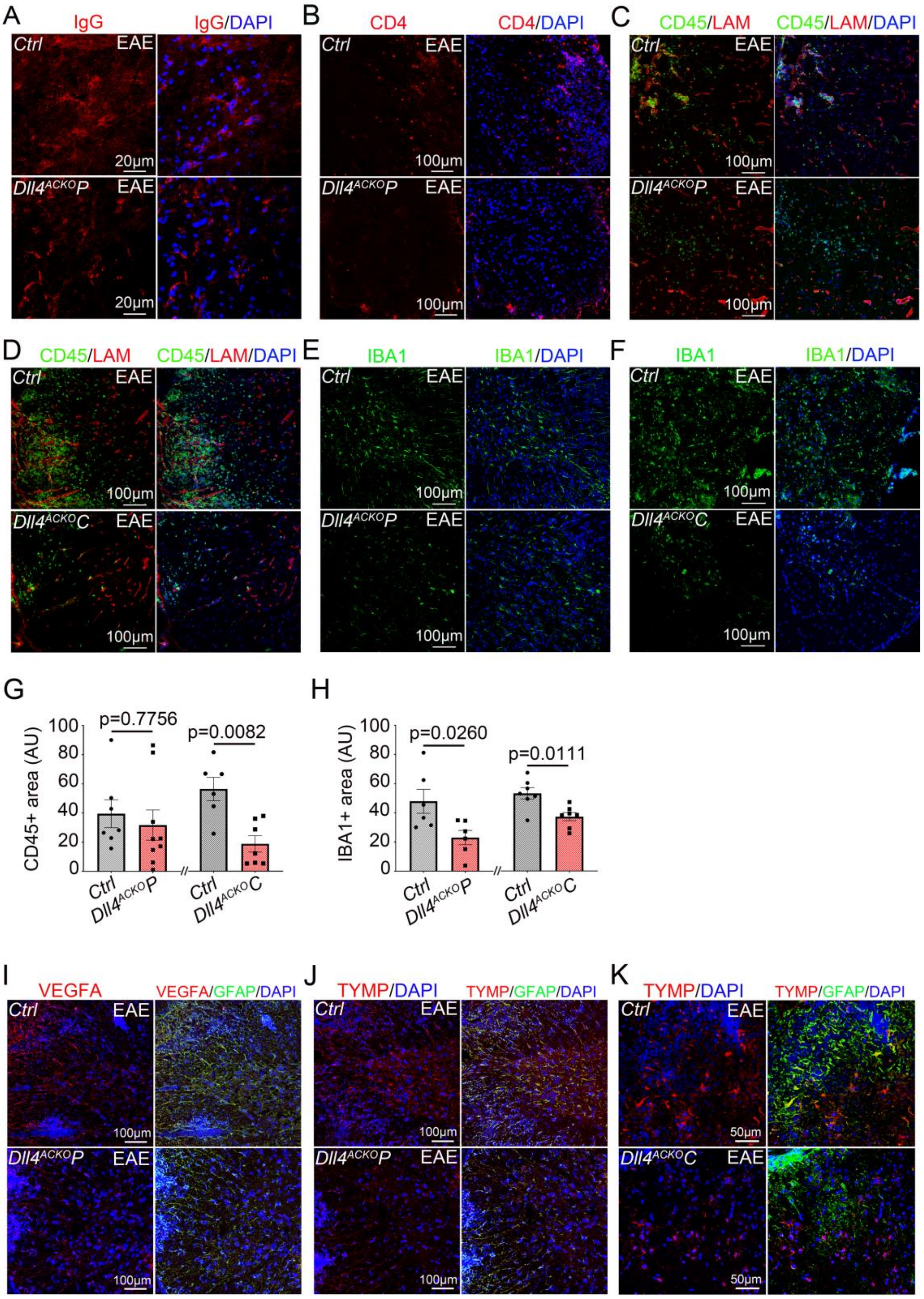


**Supplemental Fig 3 (Related to Fig 6): *Dll4$^{ACKO}$* mice display less inflammatory infiltration under EAE condition. (A-B)** Spinal cord EAE lesions from *Dll4$^{ACKO}$P* mice and littermate *controls* were harvested at 18 days post induction. *Dll4$^{ACKO}$P* and *control* lesions were immune-stained with **(A)** an anti IgG (in red) antibody and **(B)** an anti CD4 (in red) antibody. Nuclei were stained with DAPI (in blue). **(C-F):** Spinal cord EAE lesions from *Dll4$^{ACKO}$C mice, Dll4$^{ACKO}$P* mice and littermate *controls* were harvested at 18 days post induction. **(C-F)** *Dll4$^{ACKO}$P* and *control* lesions were immune-stained with **(C-D)** anti-CD45 (in green) and anti-LAM (in red) antibodies and **(E-F)** an anti IBA1 (in green) antibody. Nuclei were stained with DAPI (in blue). **(G-H)** CD45 and IBA1 positive areas were quantified (*Dll4$^{ACKO}$P* mice n = 6-7, WT n = 6-7) (*Dll4$^{ACKO}$C* mice n = 7, WT n = 6-7).

**Supplemental Fig 3 (Related to Fig 6): VEGFA and TYMP are downregulated in *Dll4$^{ACKO}$P* mice and TYMP is downregulated in *Dll4$^{ACKO}$C* mice when compared to *controls*. (I-J)** Spinal cord EAE lesions from *Dll4$^{ACKO}$P* mice and littermate *controls* were harvested at 18 days post induction. *Dll4$^{ACKO}$P* and *control* lesions were immune-stained with anti-GFAP (in green) and **(I)** anti-VEGFA (in red) or **(J)** anti-TYMP (in red) antibodies. Nuclei were stained with DAPI (in blue). **(K)** Spinal cord EAE lesions from *Dll4$^{ACKO}$C* mice and littermate *controls* were harvested at 18 days post induction. *Dll4$^{ACKO}$C* and *control* lesions were immune-stained with an anti-GFAP (in green) and anti-TYMP (in red) antibodies. Nuclei were stained with DAPI (in blue). Mann-Whitney U test.

**Supplemental Fig 4: Conditional astrocyte *Dll4* inactivation reduces the size of inflammatory CNS lesions in a model of acute neuroinflammation: (A-L)** Cerebral cortices of 10-week-old

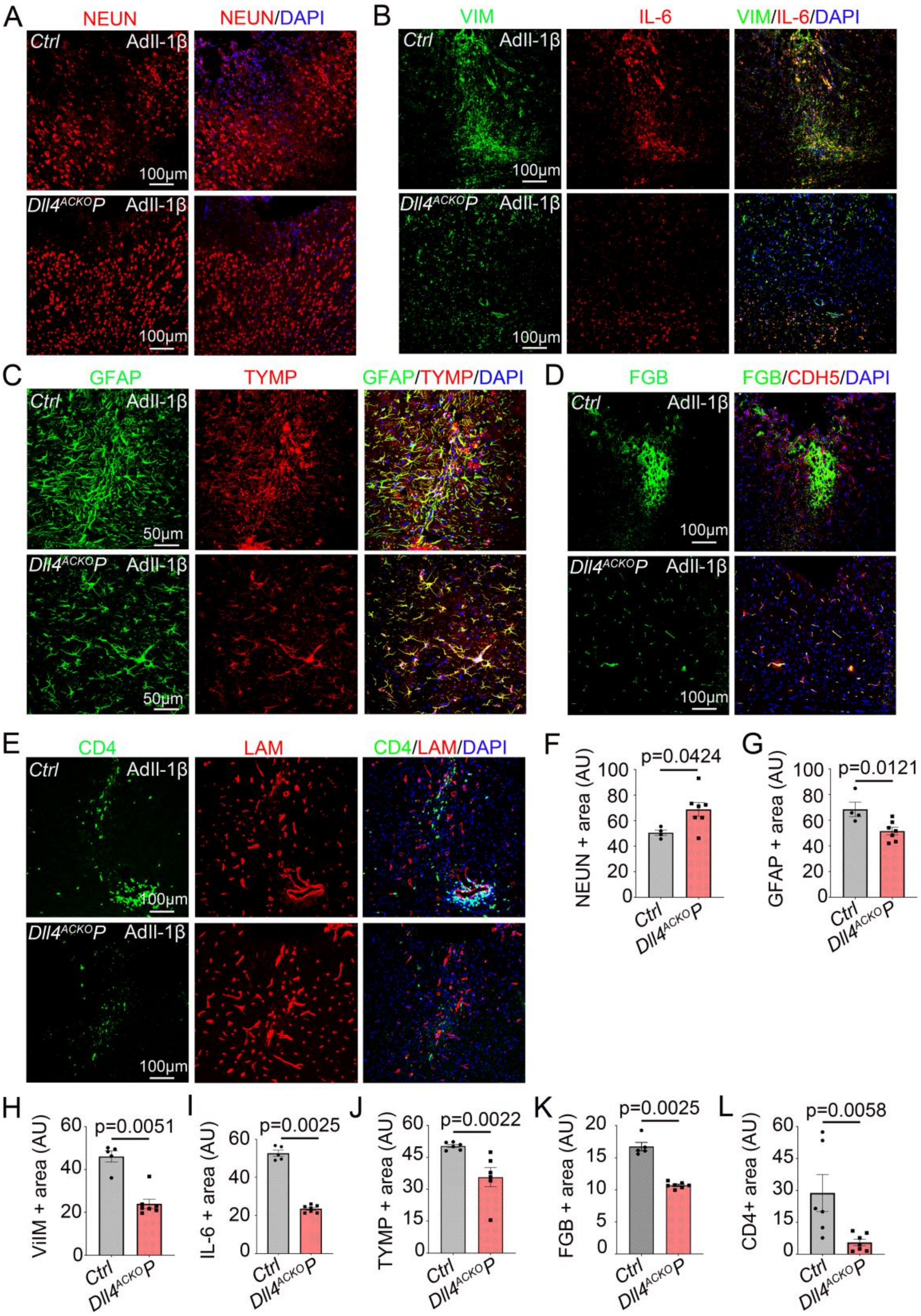


*Dll4*$^{ACKO}$*P* mice and *control* littermates were harvested 7 days following stereotactic microinjection of AdIL-1β ($10^7$ PFU). Cortical lesions were immuno-stained with **(A)** an anti-NEUN antibody, **(B)** anti-VIM (in red) and anti-IL-6 (in green) antibodies, **(C)** anti-GFAP (in green) and anti-TYMP (in red) antibodies, **(D)** anti-FGB (in green) and anti-CDH5 (in red) antibodies, and **(E)** anti-CD4 (in green) and anti-LAM (in red) antibodies. **(A-E)** Nuclei were stained with DAPI (in blue). **(F)** NEUN, **(G)** GFAP, **(H)** VIM, **(I)** IL-6, **(J)** TYMP, **(K)** FGB and **(L)** CD4 positive areas were quantified (*Dll4*$^{ACKO}$*P* mice $n$ = 6 to 7, WT $n$ = 5 to 6). Mann-Whitney U test.

**Supplemental Fig 5 (Related to Fig 8): Mice with endothelial specific *Dll4* inactivation display equivalent disability to *control* littermates in a model of multiple sclerosis: (A)** *Cdh5-Cre*$^{ERT2}$, *Dll4*$^{Flox/Flox}$ mice and *control* mice induced with EAE were scored daily according to a widely-used 5-point scale (EAE scoring: 1 limp tail; 2 limp tail and weakness of hind limb; 3 limp tail and complete paralysis of hind legs; 4 limp tail, complete hind leg and partial front leg paralysis), nonlinear regression (Boltzmann sigmoidal). (*Cdh5-Cre*$^{ERT2}$, *Dll4*$^{Flox/Flox}$ $n$ = 14, WT $n$ = 15). **(B)** *Cdh5-Cre*$^{ERT2}$, *Dll4*$^{Flox/Flox}$ and *control* mice EAE peak score and **(C)** EAE average score during time of disability were quantified other the course of the disease.

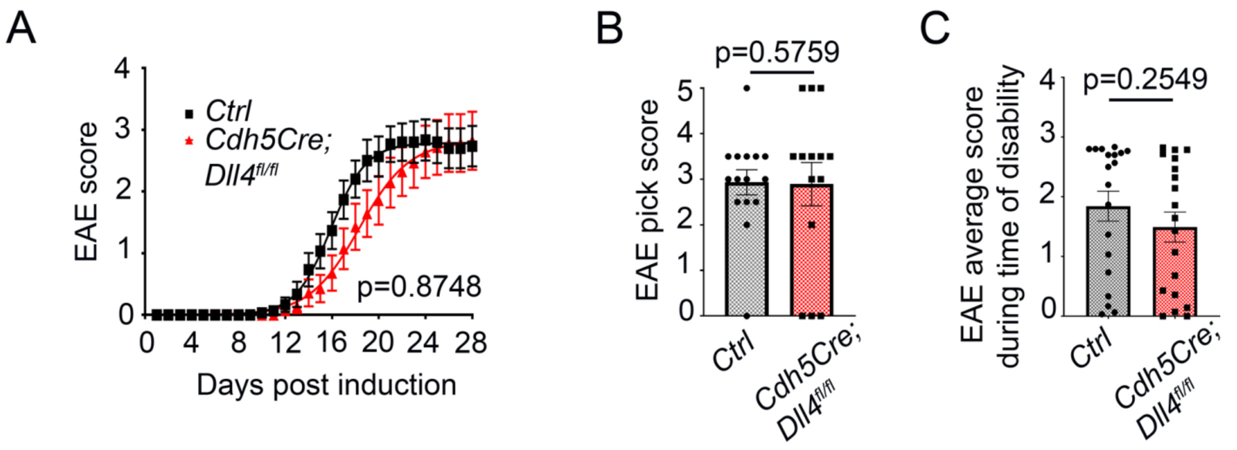
